# A multi-data based quantitative model for the neurovascular coupling in the brain

**DOI:** 10.1101/2021.03.25.437053

**Authors:** Sebastian Sten, Henrik Podéus, Nicolas Sundqvist, Fredrik Elinder, Maria Engström, Gunnar Cedersund

## Abstract

The neurovascular coupling (NVC) forms the foundation for functional imaging techniques of the brain, since NVC connects neural activity with observable hemodynamic changes. Many aspects of the NVC have been studied both experimentally and with mathematical models: various combinations of blood volume and flow, electrical activity, oxygen saturation measures, blood oxygenation level-dependent (BOLD) response, and optogenetics have been measured and modeled in rodents, primates, or humans. We now present a first inter-connected mathematical model that describes all such data types simultaneously. The model can predict independent validation data not used for training. Using simulations, we show for example how complex bimodal behaviors appear upon stimulation. These simulations thus demonstrate how our new quantitative model, incorporating most of the core aspects of the NVC, can be used to mechanistically explain each of its constituent datasets.

## 1 Introduction

The adult brain is an energy-costly organ that consumes ∼20 % of a human’s total energy consumption [1]. Despite this high energy demand, the brain constitutes only ∼2 % of the total bodyweight of an average adult [1]. To meet the energy demand, the brain requires a continuous supply of metabolic substrates such as glucose and oxygen [2]. This continuous supply of substrates is met by diffusion and receptor-mediated transport from the blood to cerebral tissue through the capillaries. Thus, as first reported by Roy and Sherrington in 1890 [3], there is a tight temporal and spatial connection between brain activity and hemodynamics. This connection is commonly referred to as the neurovascular coupling (NVC). The NVC describes the coupling between neural cells and blood vessels and involves changes in the cerebral blood flow (CBF), cerebral blood volume (CBV), and the cerebral metabolic rate of oxygen (CMRO_2_). The NVC is mediated by the synaptic activity of neurons – brain activity – that trigger the release of different vasoactive molecules [4–8] (see Fig. 1A for an overview of central signaling pathways). These molecules induce changes in CBF and CBV by acting on vascular smooth muscle cells and pericytes, which enwrap arterioles and capillaries, respectively. NVC is important because it underpins neuroimaging techniques that are based on the usage of hemodynamic changes as a proxy for neural activity [9, 10], such as the widely used functional magnetic resonance imaging (fMRI) [11]. However, the interpretation of fMRI data is usually done in a statistical way that ignores the understanding of the underlying biological mechanisms. This is because a quantitative and useful mechanistically resolved model for the NVC remains to be determined.

**Figure 1.**
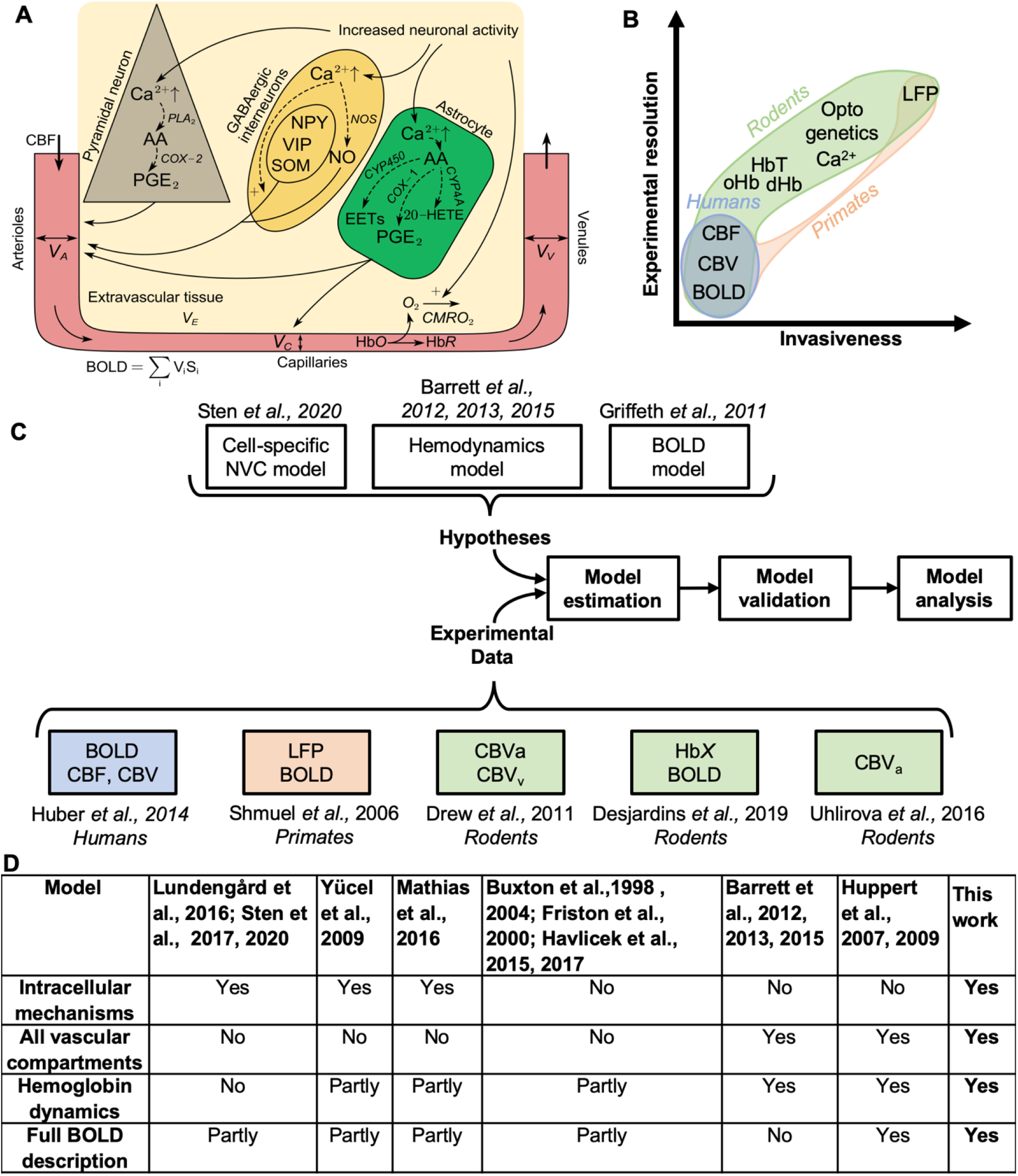
Overview of the presented study. A: Simplified schematic illustration of the cellular pathways underlying the NVC. Neuronal signaling activates GABAergic interneurons, pyramidal neurons, and astrocytes by stimulating the influx of Ca^2+^, and increases the cerebral metabolic rate of oxygen (CMRO_2_). Ca^2+^ facilitates signaling pathways in the different cells. In GABAergic interneurons, Ca^2+^ promotes nitric oxide (NO) through the upregulation of nitric oxide synthase (NOS). Furthermore, Ca^2+^ also facilitates the release of different neuropeptides: neuropeptide Y (NPY), vasoactive intestinal peptide (VIP), and somatostatin (SOM). In pyramidal neurons, Ca^2+^ promotes the synthesis of the arachidonic acid (AA) via phospholipase A2 (PLA_2_), which in turn is metabolized to prostaglandin E2 (PGE_2_) via cyclooxygenase-2 (COX-2). In astrocytes, arachidonic acid is metabolized into three different vasoactive messengers: PGE_2_ via cyclooxygenase-1 (COX-1), epoxyeicosatrienoic acid (EET) via cytochrome P450 (CYP) epoxygenase, and 20-Hydroxyeicosatetraenoic acid (20-HETE) via CYP4A. Together these vasoactive messengers act on arterioles and capillaries to modulate the vessel diameters, inducing changes in blood flow and volume. These flow and volume changes are propagated throughout the different vessels, giving rise to changes in oxygen saturation, hemoglobin concentrations, and finally, the BOLD signal. **B: Overview of commonly used experimental techniques, depicted along two axes**: the spatial and temporal resolution of what is measured, and how invasive the technique is. **C: Overview of the study**. Already published experimental data from different species and/or experimental techniques are combined with the development of a mathematical model, which builds upon three already published models. **D:** Comparison of different published models describing the neurovascular coupling (NVC) with regards to different mechanisms. The notations used in the table is: ‘Yes’ if the model features the stated mechanism, ‘Partly’ if the model has a description that is not fully satisfying, and ‘No’ if there is no description of the mechanism. As seen, no model exists that can describe every mechanism. HbX: hemoglobin measures; CBV_a_, CBV_v_: compartment specific measures of CBV.

Previous studies of the NVC have employed a variety of different measurement strategies, each with their own possibilities, strengths, and drawbacks [12]. One technique is the usage of *in vitro* experiments where brain slices are studied to elucidate intracellular signaling intermediaries in different cell types ([5, 6], and references therein). While this technique allows for detailed probing of intracellular mechanisms, it does not allow for probing the interaction between these cell types and blood flow. Another technique is invasive *in vivo* animal experiments. Such experiments are predominantly carried out during anesthesia in rodents [13, 14]. While such experiments have limited capability to measure intracellular signaling intermediaries, they allow for examinations of the crosstalk between blood supply and brain activity, and for measurements of a wide variety of entities, such as CBF, CBV, vessel diameter, hemoglobin and oxygen saturation changes, as well as electrical activity such as local field potential (LFP) [13–15]. A key possibility in such animal experiments is the usage of optogenetic techniques [16], which allows for the activation of specific neuronal cell types. While most such animal experiments are done in rodents, Logothetis *et al*., have done important studies in higher primates [9, 10, 17, 18]. In two of these studies, they measured LFP and fMRI simultaneously [9, 10]. Most of these animal studies are performed with the use of anesthetics, which affects neuronal excitability, cerebral metabolism, vascular reactivity amongst other physiological processes [19–22], which complicate the interpretation of such studies. Finally, a variety of non-invasive techniques are also available in humans, most of which use magnetic resonance imaging (MRI). Using MRI, one can measure blood oxygen level-dependent (BOLD) responses to different stimuli, and also measure CBF and CBV. In summary, components of the NVC can be measured by different techniques in different experimental systems, but no system alone allows for simultaneous measurement of all these components (see Fig. 1B for a condensed overview), and especially not in humans.

The NVC is highly conserved across different species, but despite this, there does not exist a systematic overview or synthesis of different data sources. The NVC is often non-invasively characterized using the canonical hemodynamic response function (HRF) extracted from BOLD-fMRI. The HRF consists of two or three different phases: a debated initial dip, the main response, and a post-peak undershoot. The shape of the HRF is consistent across species, such as rodents, primates, and humans. Furthermore, the timing is preserved across these species: in response to a sub-second stimulus, the peak of the main response lies at approximately three to six seconds after a stimulus, and the entire HRF usually lasts 15-20 seconds [23–25]. Because of this high conservation across species, it is relevant to consider data from different sources together, since they all provide different pieces to the same puzzle. One way to lay such a systems-level puzzle considering data from different experimental systems is to use mathematical modeling.

Previously, various aspects of the NVC have been modeled. In the standard interpretation of BOLD-fMRI, activity is equated by correlation with the canonical HRF ([26] and references therein). This approach can provide an estimate of where, in the brain, neuronal activity is present, but this approach ignores the neural and vascular mechanisms involved in the NVC. The first serious attempt at describing these mechanisms was made with the ‘Balloon’ model [27–29]. In this model, the volume of the venules is described by similar mechanisms as those governing the expansion of a balloon. While the Balloon model describes a reasonable series of steps involving neuronal activity, CBF, CBV, CMRO_2_, and the BOLD signal, the equations describing each of these individual steps are purely phenomenological. The most mechanistically correct way to describe blood flow in vessels is to solve the Navier-Stokes equations [30, 31]. In practice, a simplified approach called Windkessel models using lumped parameter models and ordinary differential equations is typically sufficient to provide a sufficient description of the interplay between CBF and CBV [32, 33]. There are a few models that describe the interplay between Windkessel dynamics in arterioles and venules, and their interplay with the BOLD signal [28, 34–37]. However, no such model describes the crosstalk between these processes and the intracellular signaling resulting in the release of vasoactive substances (Fig. 1A). Nevertheless, there are a few models that describe these intracellular signaling cascades and their impact on the blood flow regulation, even though these models use a single blood compartment [38– 40]. A summary of previous NVC models is found in Fig. 1D. One of the more advanced models describing intracellular pathways is based on optogenetics data, which has unraveled the crosstalk between pyramidal cells and two different types of interneurons [41]. However, no model of the NVC describes such intracellular pathways and all of the above described data available in rodents, primates, and humans.

Herein, we present a first mathematical model that covers multiple aspects of the NVC into one complete model (Fig. 1C & 2A–C). We have connected a mechanistic NVC model for the control of arterioles (presented in [41]) with a Windkessel model of blood flow, pressure, volume, and hemoglobin content in arterioles, capillaries, and venules (presented by Barrett *et al*. [34, 42, 43]), which finally is combined into a complete description of the BOLD signal using the model by Griffeth *et al*. [44]. Our approach is the first model that is able to describe previously published data from different species (humans, mice, and macaques), consisting of multiple different measurement variables (diameter change of the vessels, different measures of CBV, CBF, transient hemoglobin changes, BOLD responses, and LFP) gathered using both optogenetic and sensory perturbations. Furthermore, the model can successfully predict independent experimental data used for validation. Using the model, we provide mechanistic insights explaining the dynamic behavior seen in each dataset by illustrating how this model can be used to unravel the mechanisms of complicated dynamics observed in different available data sources. To the best of our knowledge, this model constitutes the most complete mathematical description of the NVC to date.

## 2 Methods

### 2.1 Model Formulation

The model is formulated using a mixture of ordinary differential equations (ODEs) and differential algebraic equations (DAEs). The general notation of such a model is as follows:

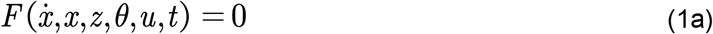

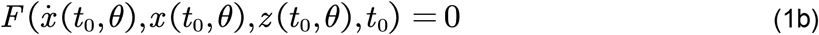

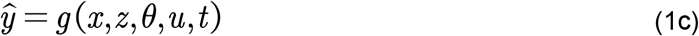

in which *x* is the vector of state variables for which derivatives are present; *z* is the vector of state variables with no specified derivatives; 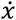 represent the derivatives of the states with respect to time *t* ; *F* and *g* are non-linear smooth functions; *θ*is the vector of unknown constant parameters; where *u* is the input signal corresponding to experimental data; where eq. 1b is the initial value solution, solving the values of the states at the initial time point *t* =*t* _0_, which are dependent on the parameters *θ*; and where *ŷ* are the simulated model outputs corresponding to the measured experimental signals. Note that 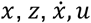 and *ŷ* depend on *t*, but that the notation is dropped unless explicitly needed.

### 2.2 Model structure

The model interaction graph is depicted in Figure 2. The interaction graph describes all of the reactions and interactions of the model. In practice, the presented model is the combination of multiple previous studies, where Section 2.2.1–2.2.3 builds upon the work of [41], 2.2.4–2.2.5 corresponds to the work of Barrett *et al*. [34, 42, 43], and finally, 2.2.6 corresponds to the work of Griffeth *et al*. [44].

**Figure 2.**
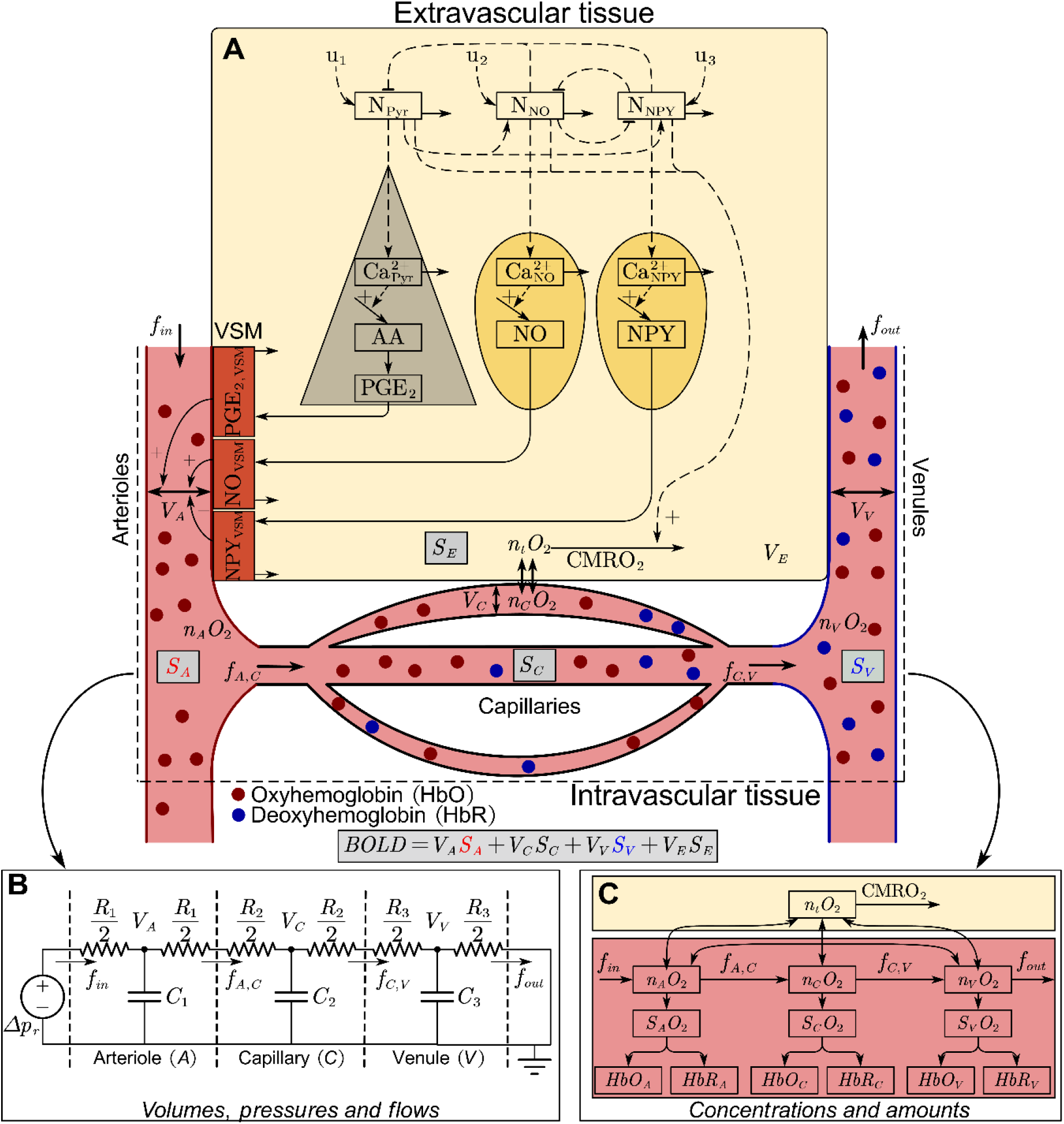
Interaction graph of the model. A: A simplified overview of the model described in [41]. Two distinct types of cells are described: pyramidal neurons (grey triangle, left) and GABAergic interneurons (divided into NO- and NPY-expressing neurons; yellow oval shapes, right). These neurons are connected in a simple relationship, with pyramidal neurons exciting GABAergic neurons, which in turn inhibit the surrounding cells. Activation of a neuron causes an influx of Ca^2+^ ions. In pyramidal neurons, Ca^2+^ activates phospholipases, which converts phospholipids into arachidonic acid (AA). AA is further metabolized into prostaglandin E2 (PGE_2_). In NO-expressing interneurons, Ca^2+^ activates nitric oxide synthase, which triggers the production and release of nitric oxide (NO). In other subtypes of GABAergic interneurons, vesicle-bound vasoactive peptides such as neuropeptide Y (NPY) are expressed. The release of these peptides is facilitated by Ca^2+^. Vascular smooth muscle (VSM) cells (brown rectangles) enwrap arterioles (left side of the vascular tree) and regulates the arteriolar diameter. PGE_2_ promotes arteriolar dilation by activation of the prostaglandin EP_4_ receptor located on the VSM. NO diffuse freely over cellular membranes, and acts to increase the production of cyclic guanosine monophosphate, which in turn promotes arteriolar dilation. Lastly, NPY activates the G-protein coupled NPY Y1 receptor expressed on VSM cells, promoting arteriolar constriction. These three effects on the VSM control the arteriolar diameter and thereby also the volume and flow of blood through the arterioles. These volume and flow changes are propagated through the capillaries and venules. **B: Three-compartment vascular model of blood volumes, pressures, and flows corresponding to an analog electrical circuit, as described in [34]**. The blood pressure drop corresponds to a voltage drop, the blood flow to electric current, and the blood volume to electric charge stored in the capacitors. The vessel compliance plays the role of capacitance, and the vessel resistance is analogous to electric resistance. The blood pressure difference maintained by the circulatory system corresponds to the electromotive force. **C: Oxygen transport model, as described in [42]**. The diagram depicts amount of oxygen (n_i_O_2_), oxygen saturation (S_i_O_2_), oxygenated hemoglobin (HbO_i_) and deoxygenated hemoglobin (HbR_i_), for each respective compartment (A = arteriole/arterial, C = capillary, V = venule/venous, t = tissue). Oxygen in tissue can be metabolized, indicated by the cerebral metabolism of O_2_ (CMRO_2_) arrow leaving the state. All of these different models in unison affect the blood oxygenation and blood volume in each respective compartment, which in turn determines the specific compartments contribution (S_i_) to the BOLD-fMRI signal (grey boxes).

**Figure 3.**
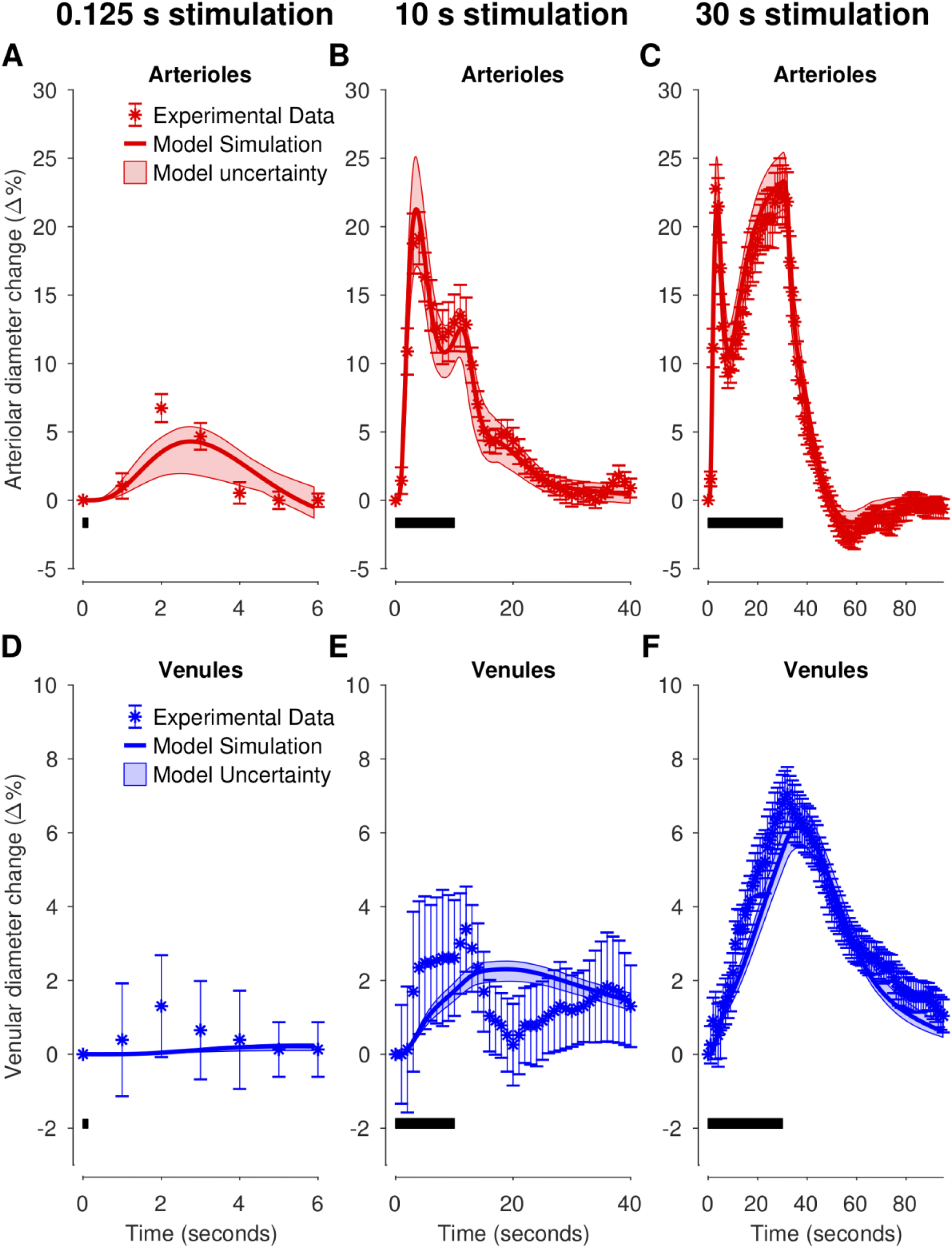
Model estimation to experimental data of arteriolar (A–C) and venular (D–F) diameter changes in awake mice for three different sensory stimulation lengths: 125 ms (**A & D**), 10 s (**B & E**), and 30 s (**C & F**). Experimental data are replotted versions of data presented in Fig. 2C of the original manuscript by Drew *et al*. [75]. The stimulation lengths are denoted with the black bar in the bottom left portion of each graph. For each graph: experimental data (colored symbols); the uncertainty of the experimental data is presented as SEM (colored error bars); the best model simulation is seen as a colored solid line; the model uncertainty as a colored semi-transparent overlay. The x-axis represents time in seconds, and the y-axis is the normalized vessel diameter change (Δ%).

The stimulus *u* is given by the following equation:

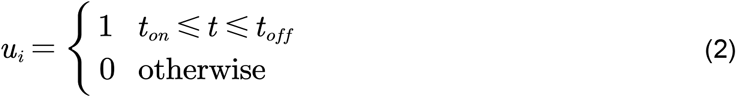

where *t* _*on*_, *t* _*off*_ are the times when the signal goes on and off, respectively.

#### 2.2.1 Presynaptic activity and calcium influx

Glutamate and GABA are released from different types of neurons upon membrane depolarization and bind to specific ion channel-coupled receptors located in the neuronal plasma membrane. Glutamate, an excitatory neurotransmitter, binds to *α*-amino-3-hydroxy-5-methyl-4-isoxazolepropionic acid (AMPA) and *N*-methyl-D-aspartate (NMDA) receptors. Activation of these ion channel-coupled receptors triggers ion-conducting pores to open, triggering an influx of Na^+^ and Ca^2+^ ions which cause a depolarization of the neuron. Depolarization opens voltage-gated calcium channels, allowing a further influx of Ca^2+^ ions. In contrast, *γ*-aminobutyric acid (GABA) acts to prevent depolarization of neurons as it binds to the ion channel-coupled GABA_A_ or GABA_B_ receptor, which opens Cl^-^ ion-conducting pores. Pyramidal neurons are glutamatergic, meaning that glutamate is released upon depolarization, and acts on *e*.*g*., astrocytes and interneurons. GABAergic interneurons release GABA upon depolarization and target pyramidal neurons or other interneurons. This forms a simple relationship, where interneurons regulate the activity of pyramidal neurons. In the model, we represent these interplays between pyramidal and inhibitory interneuron activity and their combined effect on respective neuronal Ca^2+^ levels as

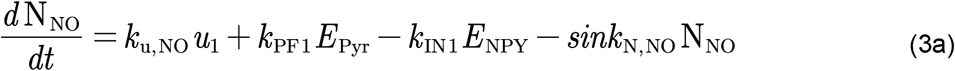

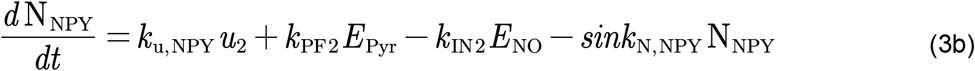

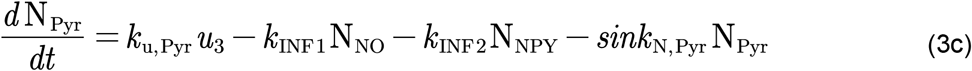

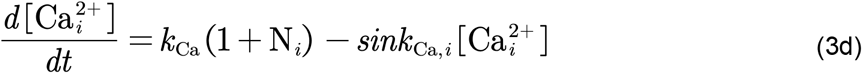

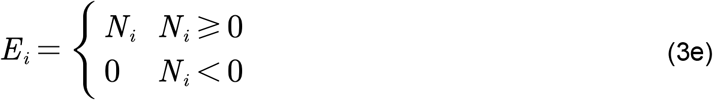

Where N_NO_, N_NPY_, and N_Pyr_ describe the neuronal activity of the GABAergic nitric oxide (NO) and neuropeptide Y (NPY) interneurons, and the pyramidal neurons, respectively; where 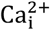 represents intracellular calcium in each respective neuron; *k*_u,i_ is the input strength of the stimulation to respective neuron; where *k*_PF1_ and *k*_PF2_ are the kinetic rate parameters governing pyramidal to GABAergic interneuron signaling; where *k*_INF1_ and *k*_INF2_ are the kinetic rate parameters describing the negative feedback from GABAergic interneurons to pyramidal neurons; where *k*_IN1_ and *k*_IN2_ are the kinetic rate parameters describing the negative feedback between GABAergic interneurons; where *sink*_N,i_ are kinetic rate parameters governing the degradation of the activity of respective neuron; where *k*_Ca_ is the basal inflow rate of Ca^2+^ (*k*_Ca=_10 for all three neurons), and finally, where *sink*_Ca,i_ is the elimination rate of intracellular Ca^2+^, and i = (Pyr, NO, NPY) denotes the three different neuronal types.

#### 2.2.2 Pyramidal neuron signaling

The rise in intracellular Ca^2+^ levels in pyramidal neurons activates phospholipases which metabolize, through intermediary enzymatic steps, membrane phospholipids into intracellular arachidonic acid (AA) [12, 45, 46]. This is described by:

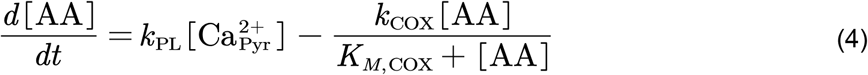

where *k*_PL_ and *k*_COX_ are kinetic rate parameters and *K*_*M,COX*_ is a Michaelis constant. In pyramidal neurons, AA is metabolized into prostaglandin E_2_ (PGE_2_) through a cyclooxygenase-2 (COX-2) and PGE synthase rate-limiting reaction [45, 47]. PGE_2_ evoke vasodilation by through activation of EP2 and EP4 receptors expressed on the surface of vascular smooth muscle cells (VSM) cells [45, 48]. In the model, this mechanism described as

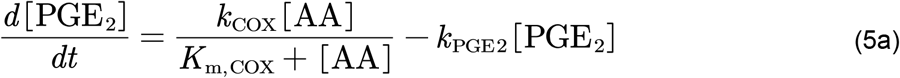

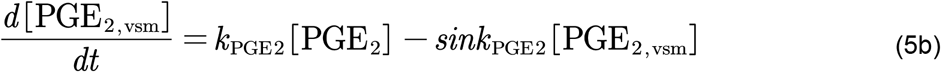

Where PGE_2,vsm_ represents PGE_2_ acting on the VSM cells, and *k*_PGE2_ and *sink*_PGE2_ are kinetic rate parameters.

#### 2.2.3 GABAergic interneuron signaling

The rise in intracellular Ca^2+^ levels in GABAergic interneurons evoke the release of different vasoactive messengers and substances. For instance, NO has previously been shown to be a potent vasodilator at the level of arteries and arterioles in both *in vitro* and *in vivo* studies [49–52]. NO is released by specific NO-interneurons through a Ca^2+^ dependent nitric oxide synthase (NOS) rate-limiting reaction (see [6] and references therein). Here, we represent this process as

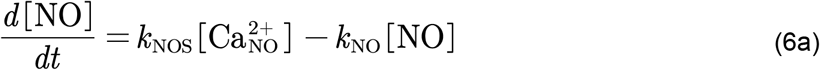

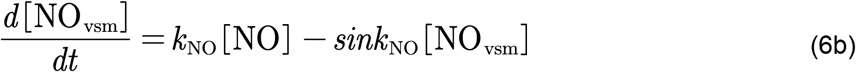

where NO_vsm_ represents NO acting on the VSM cells, and *k*_NOS_, *k*_NO_ and *sink*_NO_ are kinetic rate parameters. Here, *sink*_NO_ > 1 *s*^−1^ following experimental work by [53].

Another potent vasoactive messenger is NPY. NPY is released by specific NPY producing subtypes of GABAergic interneurons and has previously been shown to induce vasoconstriction of vessels *in vitro* [49, 54] and *in vivo* [55]. NPY binds to the NPY receptor Y1, a G_αi_-protein coupled receptor expressed on the surface of VSM cells enwrapping arteries and arterioles [56–58]. Activation of this receptor inhibits the synthesis of adenosine monophosphate (cAMP) and increase the intracellular Ca^2+^ [59], leading to VSM contraction and vasoconstriction. In the model, these mechanisms are represented as:

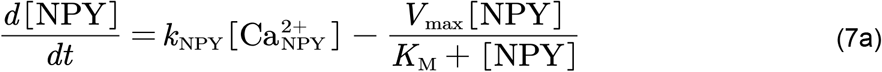

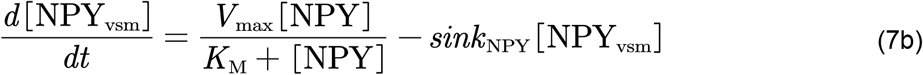

where NPY_vsm_ represents NPY acting on the VSM cells, and *k*_NPY_ and *sink*_NPY_ are kinetic rate parameters. The conversion between intracellular NPY and NPY in the VSM, NPY_vsm_, is governed by Michaelis-Menten kinetics [60, 61], where *K*_*M*_ is the Michaelis constant (the concentration of the substrate at which half max reaction rate is achieved) and V_max_ is the maximal reaction rate.

The expression of the different vasoactive substances is scaled with a parameter, 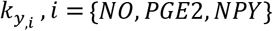, and summarized to a total vascular influence *G*:

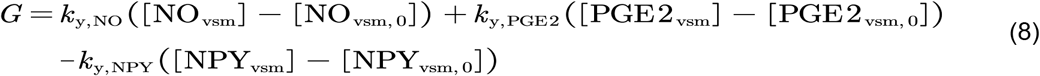

#### 2.2.4 Cerebrovascular dynamics described by an electrical circuit analog model

We describe the dynamics of three cerebrovascular compartments, corresponding to arteries/arterioles (*a*), capillaries (*c*), and veins/venules (*ν*), respectively. The compartments are represented by an electrical circuit analogy, as originally presented in the work by Barrett *et al*. [34, 42, 43]. We use the derivative work of [62], and only present the fully derived equations to improve the clarity for the reader. We refer the reader to the original articles for an in-depth description of the equations [34, 42, 43, 62]. The following equations describe the volume change for each respective cerebrovascular compartment, V_*i*_, *with i*={*a, c, ν*}.

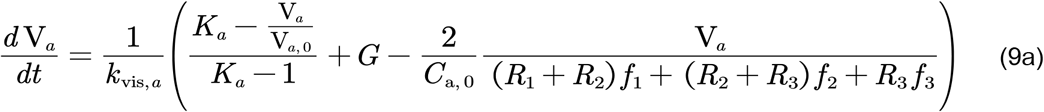

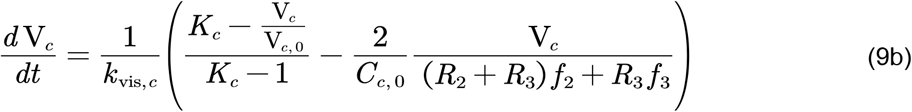

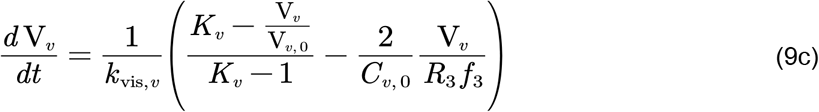

Here, *K* are stiffness coefficients; *k*_*νis,i*_ are viscoelastic parameters; *C*_*i*_ represents the compliance of the vessel; *R* represents the vessel resistance; *f* represents the flow of blood between the cerebrovascular compartments; the baseline value is indicated by the subscript 0, and finally, *G* is the vasoactive function translating the actions of the vasoactive arms (Eq. 8) into hemodynamic changes. In short, the arterial compartment is actively regulated by the vasoactive arms (Eq. 8 & 9a), and the evoked hemodynamic changes in the arterial vessels are propagated through capillary and venous compartments.

Using conservation of mass, the rate at which the blood volume change in a compartment is given by the difference between in- and outflow of blood:

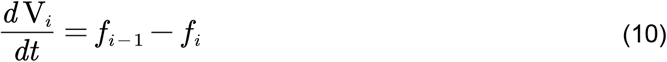

Restructuring the equation gives the following relationship:

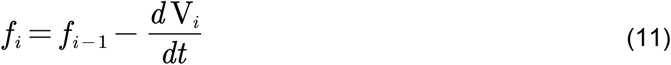

Next, the pressure drop over a compartment is given by:

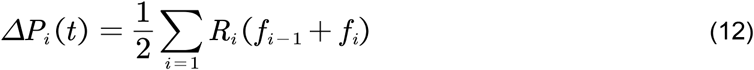

Which is subject to the pressure boundary conditions:

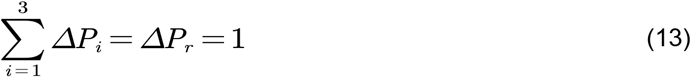

Using this conditions, we can solve for the inflow of blood into the first compartment *f*_0_:

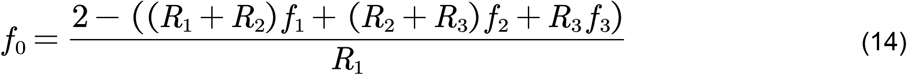

The equations above form a DAE system, comprised of three volumes and four flows. To further simplify, the scales of the variables are normalized to:

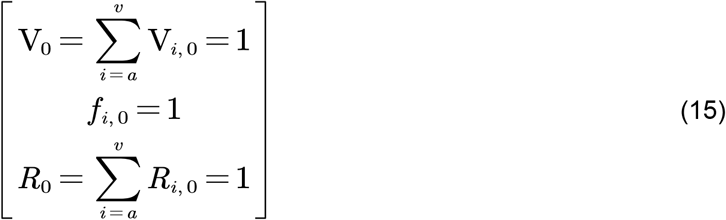

Using this, the rest of the physiological terms can be expressed as:

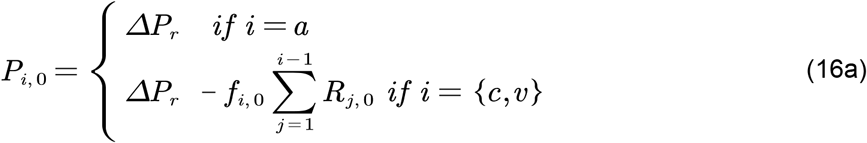

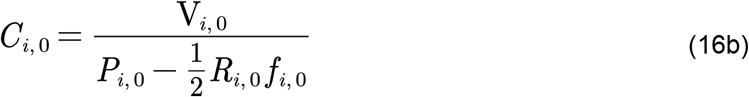

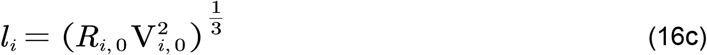

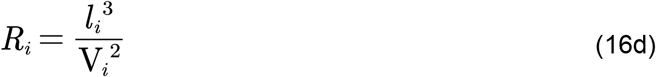

where *P* is pressure at the entry point to each compartment, *C* is the compliance of the vascular vessel, *L* is the length of the vascular segment, and *R* is the vessel resistance to flow.

Given the Equations 9–16, the initial conditions for the cerebrovascular volumes V_*i*_ and the vascular resistances *R*_*i*_ must be specified. We set these initial conditions in accordance to the work of Barrett *et al*. [34], which based these values on available literature, resulting in V_*i*,0=_[0.29, 0.44, 0.27] and *R*_*i*,0=_[0.74, 0.08, 0.18].

#### 2.2.5 Oxygen transportation

To describe the oxygen transportation through the cerebrovascular system, we describe three vascular compartments and a cerebral-tissue compartment. The amount of oxygen in these compartments is given by:

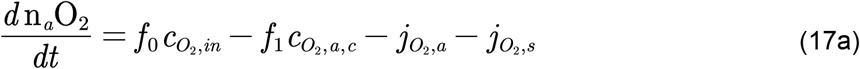

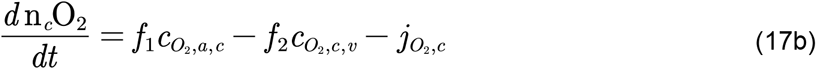

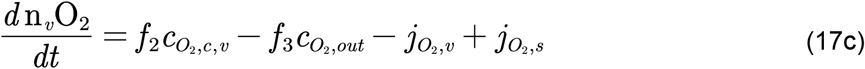

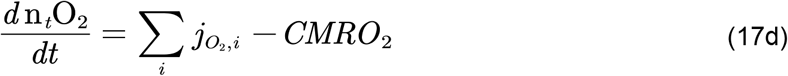

where the flow of blood *f* transports oxygen in and out from each compartment. The permeability in the vessel wall allows for oxygen to diffuse to the tissue compartment, 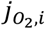, where *i* ={*a, c, ν*}. An arterio-venous diffusion shunt is described by 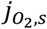, where oxygen moves from the arterial to venous compartment. Oxygen is metabolized in the tissue compartment, *CMRO*_2_, which is given by:

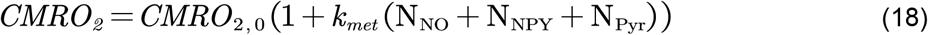

where *CMRO*_2,0_ is the basal metabolism of oxygen, which is increased by the activity-level of each neuron (see Eq. 3) scaled by the parameter *k*_*met*_.

The concentration of oxygen, 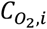 is given by the amount of oxygen, *n*_*i*_*o*_2_, divided by the volume of each compartment, V_*i*_, except for the oxygen concentration that enters the arteries, 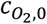 which is assumed to be constant, minus a loss of oxygen to the surrounding tissue 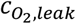.

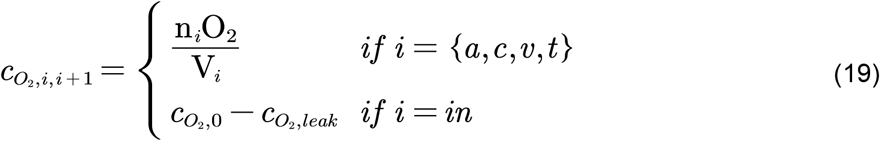

Where the tissue volume fraction V_*t* =_34.8 [42] and 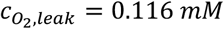 (calculated using data from [42, 63]).

As in the original work [42], by ignoring the minor fraction of oxygen directly dissolved in the blood plasma, blood oxygen concentration can be related to oxygen partial pressures through the oxygen-hemoglobin saturation curve:

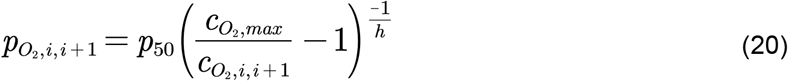

Where *p*_50=_36 *mmHg* is the oxygen partial pressure at the half-way saturation point, 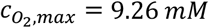 is the maximum oxygen concentration in blood, and *h*=2. 6 is the Hill exponent. These values are taken from [64].

The oxygen pressure in the tissue, 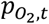 is calculated using Henry’s law:

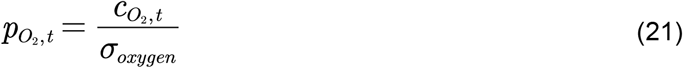

Where σ_*oxygen*_1.46 *M mmHg* is the coefficient for solubility of oxygen in tissue [65].

The average pressure 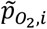 in a compartment is obtained by averaging over the input and output pressures:

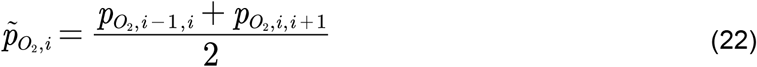

The diffusion of oxygen between the vessels and the tissue, 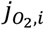 as well as between the arteries and the veins, 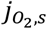, is driven by the difference in partial oxygen pressure:

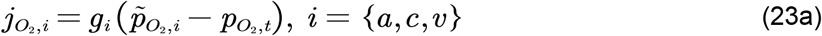

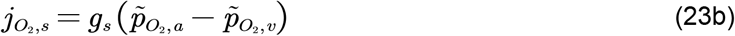

where *g*_*i*_, *i*={*a, c, ν, s*} are rate constants.

We employed the identical strategy of Barrett *et al*. [42] (see supplementary material in [42]), and used the experimental 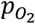 data from Vovenko *et al*. [63] to calculate *g*_*i*_, 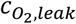 and *CMRO*_2,0_. This leaves the parameter *k*_*met*_ to be estimated where applicable.

The oxygen saturation of blood in the different vascular compartments, *S*_*i*_ *o*_2_, is approximated by dividing the average oxygen concentration of each compartment with 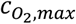:

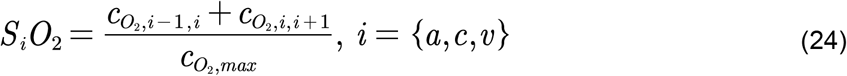

Lastly, using the blood oxygen saturation *S*_*i*_ *o*_2_ and vascular volumes V_*i*_, corresponding changes in oxygenated hemoglobin (HbO), deoxyhemoglobin (HbR), and total hemoglobin (HbT) are given by:

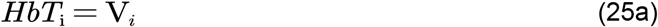

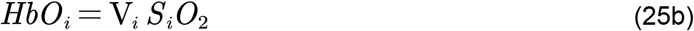

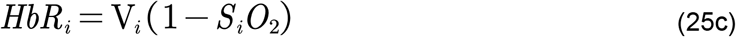

#### 2.2.6 BOLD signal derivation

Finally, the oxygen saturation of blood in the different vascular compartments, *S*_*i*_*o*_2_ (Eq. 24) and the vascular volumes, V_*i*_ (Eq. 9) are used to calculate the BOLD signal. The following derivation of the BOLD signal equations and parameter values are taken from the work of [44], and we refer the reader to the original article for a detailed view of the equations. The BOLD signal is expressed as a summation of the contributions from each vascular compartment {*a, c, ν*} and an extracellular compartment {*e*}, symbolizing the tissue:

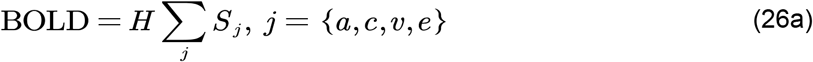

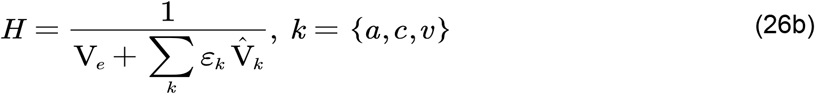

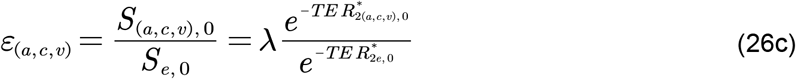

Where *S*_*j*_ is the intrinsic signal from each compartment; 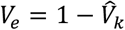 is the extravascular volume; 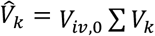 is the intravascular volume scaled by the baseline intravascular volume fraction *V*_*iν*,0_=0.05, [66]); ε_*i*_ is the intrinsic signal ratio of blood and λ is the intravascular to extravascular spin density ratio (we assume λ=1.15); 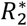 is the transverse signal relaxation rate for the four compartments, and is the echo time used during image acquisition in each study.

The transverse signal relaxation rate for the extravascular compartment 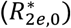 was assumed to be 25.1*s*^−1^ [67], and was calculated for the intravascular compartments by utilizing the following quadratic expression taken from [68]:

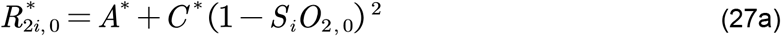

where

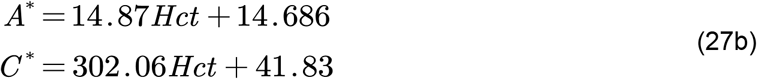

These equations are governed by *S*_*i*_*o*2 (Eq. 24) and the resting hematocrit in blood (*Hct* ; *Hct* _*a,ν*_=0.44, *Hct* _*c*=_0.33 [69, 70]).

The respective signal contribution from each compartment is given by:

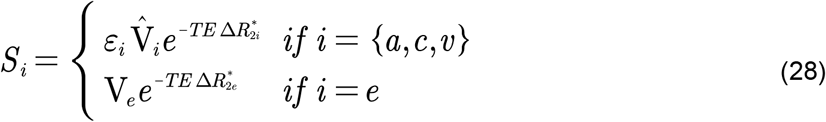

Where 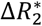 (*i*.*e*., the change in MR signal relaxation rate) is given by the following expressions:

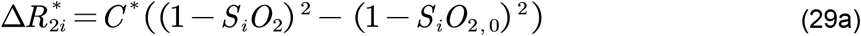

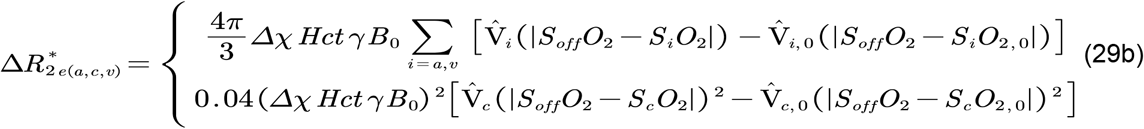

Here, Δ*χ*=2.64 × 10^−7^ is the susceptibility of fully deoxygenated blood [71]; *γ*=2.68 10^8^ is the gyromagnetic ratio of protons; *B*_0_ is the magnetic field strength used during image acquisition, and finally, *S*_*off*_*o*_2_=0.95 is the blood saturation which gives no magnetic susceptibility difference between blood and tissue [71].

### 2.3 Model evaluation

#### 2.3.1 Optimization of parameters

Once the model structure has been formulated (Fig. 2) and data has been collected (data acquisition described below), the parameters, *θ*, need to be determined. This is commonly done by evaluating the negative log-likelihood function. Assuming independent, normally distributed additive measurement noise, the likelihood of observing data given *θ*is:

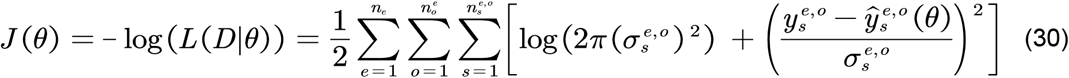

where *n*_*e*_ is the number of experiments ; where 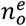 is the number of observables *o* per *e*; where 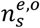 is the number of samples per *o* and *e*; where 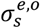 is the standard deviation of the data point; where 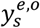 is the measured data point; where *ŷ*(*θ*)is the corresponding simulated data point. By maximizing *L*, the maximum likelihood estimate of the unknown parameters *θ* can be obtained. However, it is more common and more numerically efficient to minimize the equivalent negative log-likelihood function:

If the measurement noise 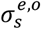 is known, *J*(*θ)* share the same optimal parameters with the least-squares function *J*_*lsq*_ :

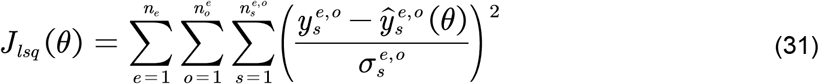

In practice, the parameters are determined by optimizing *J*(*θ*) over *θ* by using various optimization algorithms (see Section 2.5 below), and given by:

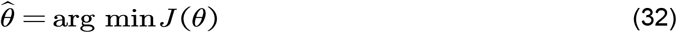

where 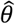 are the optimal parameters.

#### 2.3.2 Simulation uncertainty

The model uncertainty was estimated using a MCMC-sampling procedure with 10^5^ samples (see Section 2.5 for details), generating posterior distributions of the parameters, and collecting all *χ*^2^ acceptable parameters encountered. Using the acceptable *χ*^2^ parameters, a confidence interval:

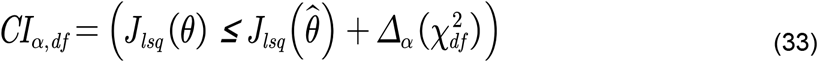

was drawn where the threshold 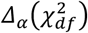 is the *α* quantile of the 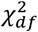 statistic [72, 73]. We also require that all acceptable parameters should pass a *χ*^2^-test.

### 2.4 Experimental data

In this work, experimental data from a total of five different studies have been used, which includes data from rodents, primates, and humans. Below, condensed descriptions of the experimental data are provided, and we kindly refer the reader to the original manuscripts for a complete description of the experimental methods. All data were extracted from figures in the respective original publications using the tool WebPlotDigitizer [74].

#### 2.4.1 Fractional diameter change of arterioles and venules in mice

The work of Drew *et al*. [75] presents experimental data of fractional diameter change in arterioles and venules upon different lengths of a sensory stimulation in mice (n = 21), measured using 2-photon imaging. The sensory stimulation consisted of trains of air puffs (delivered at 8 Hz, 20 ms duration per puff) directed at the vibrissae, with three different durations: 20 ms, 10 s, and 30 s. The mice were awake during the experiment, but were briefly anesthetized before the imaging session using isoflurane (2% in air). We have chosen to consider the period after each puff as a part of the stimulation cycle. This makes the shortest stimulation length of one air puff 125 ms long. The vascular responses were extracted from figure 2C in the original publication [75].

#### 2.4.2 Arteriolar diameter responses for OG and sensory stimuli in mice

Uhlirova *et al*. [55] reports arteriolar diameter responses to both OG and sensory stimulations measured using *in vivo* two-photon imaging, during both awake and under anesthesia conditions in mice. Briefly, four different stimulation protocols were used in their study: three conducted under the administration of the anesthetic α-chloralose, and the fourth during awake condition. Below, summarized descriptions of each stimulation protocol are presented, and we refer the reader to the original publication ([55]) and our previous modeling paper [41] for a complete description.

##### OG stimulation of pyramidal neurons during anesthesia

Thy1-ChR2-YFP mice (n = 2 subjects), with Channelrhopodsin-2 (ChR2) selectively expressed in layer 5 pyramidal neurons, were imaged (28 measurement locations along 13 arterioles within 40-380 μm range) during OG stimulation (single light pulse lasting 50-80 ms) under both control conditions and under the influence of both the AMPA/kaniate receptor antagonist 6-cyano-7-nitroquinoxaline-2,3-dione (CNQX), and the NMDA receptor-selective antagonist D-(-)-2-Amino-5-phosphonopentanoic acid (AP5).

##### OG stimulation of GABAergic interneurons during anesthesia

VGAT-ChR2(H134R)-EYFP mice (n = 3 subjects), with ChR2 selectively expressed in GABA interneurons, were imaged (52 measurement locations along 25 arterioles within 50-590 μm range) during OG stimulation (one pair of light pulses separated by 130 ms, for a total of 450 ms duration) under both a control condition and during the application of the NPY receptor Y1 inhibitor BIBP-3226.

##### Sensory stimulation during anesthesia

Additionally, four wild-type mice underwent imaging (10 measurement locations along 7 arterioles within 130-490 μm range) during sensory stimulation (2 s train of electrical pulses) under both a control condition and during the application of the NPY receptor Y1 inhibitor BIBP-3226.

##### Sensory and OG stimulation during awake condition

Lastly, four wild-type and three VGAT-ChR2(H134R)-EYFP awake mice were imaged during sensory stimulation (1 s of air puffs, 100 ms puffs at 3 Hz) and OG stimulation (single light pulse lasting 150-400 ms), respectively.

For the data extracted from this study, standard error of the mean (SEM) in all extracted time series with SEM measurements smaller than mean SEM was set to the mean SEM. This was done to avoid overfitting the model to a few extreme data points.

#### 2.4.3 Hemoglobin and BOLD responses for OG and sensory stimuli in mice

Desjardins *et al*. [76] presents experimental data consisting of optical intrinsic imaging (OIS) of blood oxygenation for both wild-type mice (n = 16) and two mice-lines transfected with (ChR2). The mice were kept in an awake state during the data acquisition. In the transfected mice, ChR2 was expressed in pyramidal neurons (Emx1-Cre/Ai32) or in all inhibitory interneurons (VGAT-ChR2(H134R)-EYFP). For the wild-type mice, a sensory stimulus consisting of air puffs (3-5 Hz) delivered to a whisker pad (duration 2 or 20 s) was used to induce hemodynamic responses. The OG-transfected mice were exposed to an OG stimulus consisting of a single 100 ms light pulse (473 nm) or a 20 s block of 100 ms light pulses delivered at 1 Hz. The blood oxygenation data were analyzed using the baseline assumption of 60 *M* HbO and 40 *M* HbR. In addition, some Emx1-Cre/Ai32 mice underwent BOLD fMRI imaging for the 20 s OG stimulus using gradient-echo echo-planar-imaging (GE-EPI) readout (TE/TR = 11–20 ms / 1 s).

#### 2.4.4 Electrophysiological and BOLD responses in macaque monkeys

The study by Shmuel *et al*. [10] reports simultaneously measured electrophysiological responses by invasive recording, and non-invasively measured BOLD responses. These measurements were carried out in the visual cortex of anesthetized macaques at 4.7 T (n=7). The stimulus paradigm consisted of a 20 s visual stimuli featuring flickering radial checkers rotating at 60°/s. A grey background persisted for 5 s before and 25 s after the stimuli. If the visual stimuli overlapped with the receptive field of V1, it induced positive BOLD responses in close vicinity to the electrode. In contrast, if the visual stimuli were presented outside the receptive field of V1, it induced negative BOLD responses in the same area. The electrophysiological recordings were processed by averaging the fractional change of the power spectrum over the frequency range of 4–3000 Hz, using a temporal resolution of 1 s. BOLD data were acquired using a gradient-echo echo-planar-imaging (GE-EPI) readout (TE = 20 ms, TR = 1 s, in-plane spatial resolution 0.75 x 0.75 x 2 mm^3^). Positive and negative BOLD responses evoked by the two visual stimuli were sampled and averaged over the same voxels located around the electrode. These processed responses were extracted from Figures 1D (BOLD) and 2A (electrophysiological responses) in the original publication [10], respectively. The SEM in all extracted electrophysiological time series with SEM measurements smaller than mean SEM was set to the mean SEM, to avoid overfitting to a few extreme datapoints.

#### 2.4.5 MR-based hemodynamic measurements in humans

The study by Huber *et al*. [77] presents *in vivo* MR-based measurements of CBF, CBV, and BOLD changes in humans (n=17) evoked using multiple visual stimuli. The study used three different visual stimulation paradigms (alternating 30 s rest vs. 30 s stimulation), consisting of two different full-field flickering checkerboards and one small circle flickering checkerboard paradigm. The full-field flickering checkerboard paradigms were used to measure arterial CBV, venous CBV, and total CBV data to both an excitatory and an inhibitory task. The small circle paradigm was used to evoke strong negative BOLD responses, and to concurrently measure both positive and negative BOLD responses, as well as the associated total CBV changes. These data were acquired using a Slice-Saturation Slab-Inversion Vascular Space Occupancy (SS-SI-VASO) sequence [78] with a multi-gradient echo EPI readout (*TE*_*1*_*/TE*_*2*_*/TE*_*3*_*/TI*_*1*_*/TI*_*2*_*/TR* = 12/32/52/1000/2000/2500/3000 ms, nominal resolution = ∼1.3 x ∼1.3 x 1.5 mm^3^). In addition, CBF changes were acquired from ROIs of positive and negative BOLD responses using a pulsed arterial spin label (PASL) sequence in four subjects (*TE*_*1*_*/TE*_*2*_*/TI*_*1*_*/TI*_*2*_*/TR* = 8.2/19.4/700/1700/3000 ms, nominal resolution = 3 x 3 x 3 mm^3^). BOLD and total CBV responses were extracted from Figure. 4A–B; arterial CBV, venous CBV, and total CBV responses were extracted from Figure. 5B–C, and finally, CBF responses were extracted Figure. 6C in the original publication.

**Figure 4.**
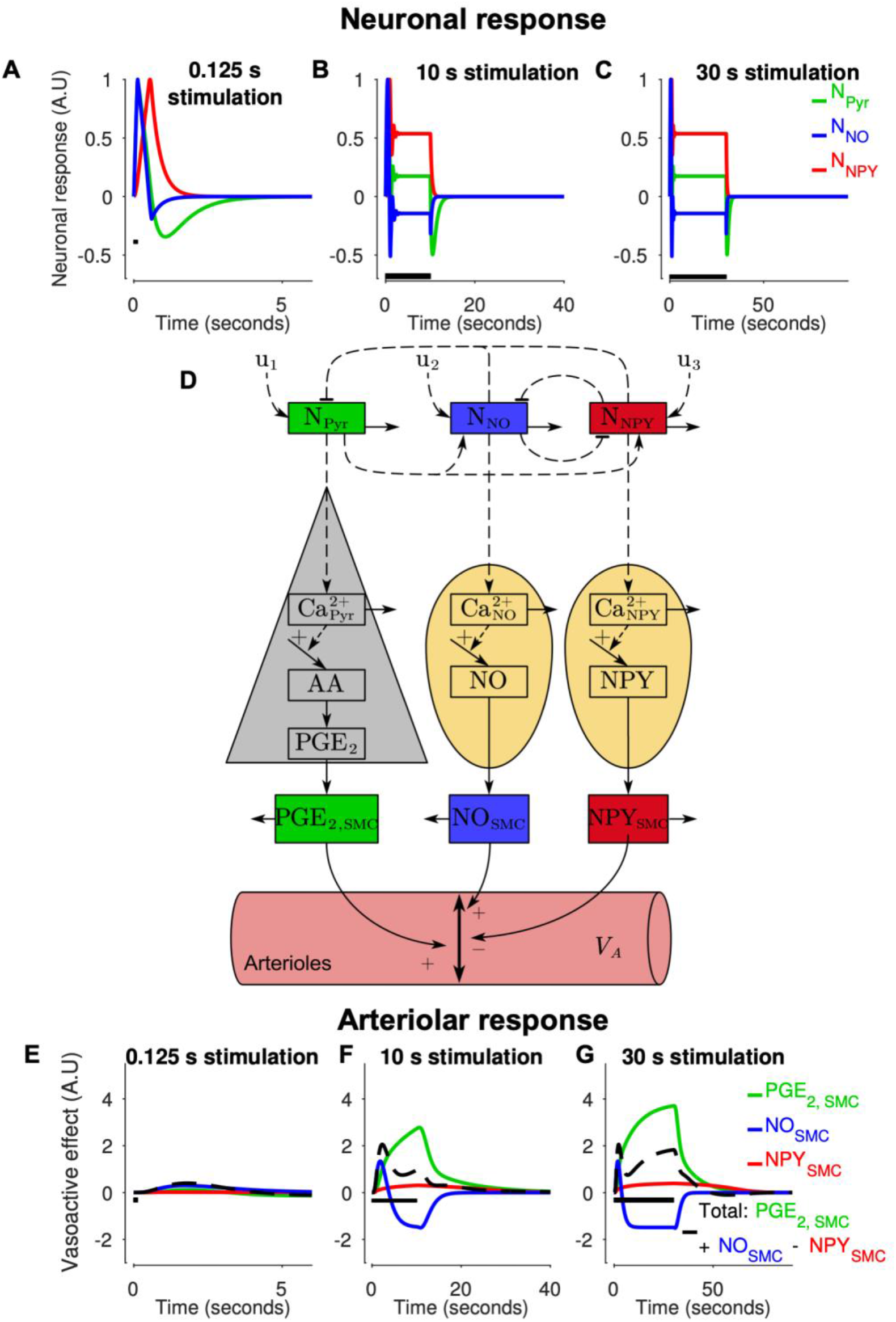
Model explanation (with the relevant model states highlighted in D) of the dynamic arteriolar behavior observed in experimental data from Drew *et al*. [75]. Model simulations for the three stimulus durations: 125 ms (**A & E**), 10 s (**B & F**), and 30 s (**C & G**) are shown. For each stimulus: the dynamic of the neuronal states N_Pyr_, N_NO_, N_NPY_ (**A–C**), and the arteriolar response (**E–G**). For each graph: model simulation (colored lines); stimulus length (black bar). The x-axis represents time in seconds, and the y-axis is the change in neuronal states (A.U) for **B–D**, and the vasoactive effect on the arteriolar compartment (A.U) for **E–G**.

**Figure 5.**
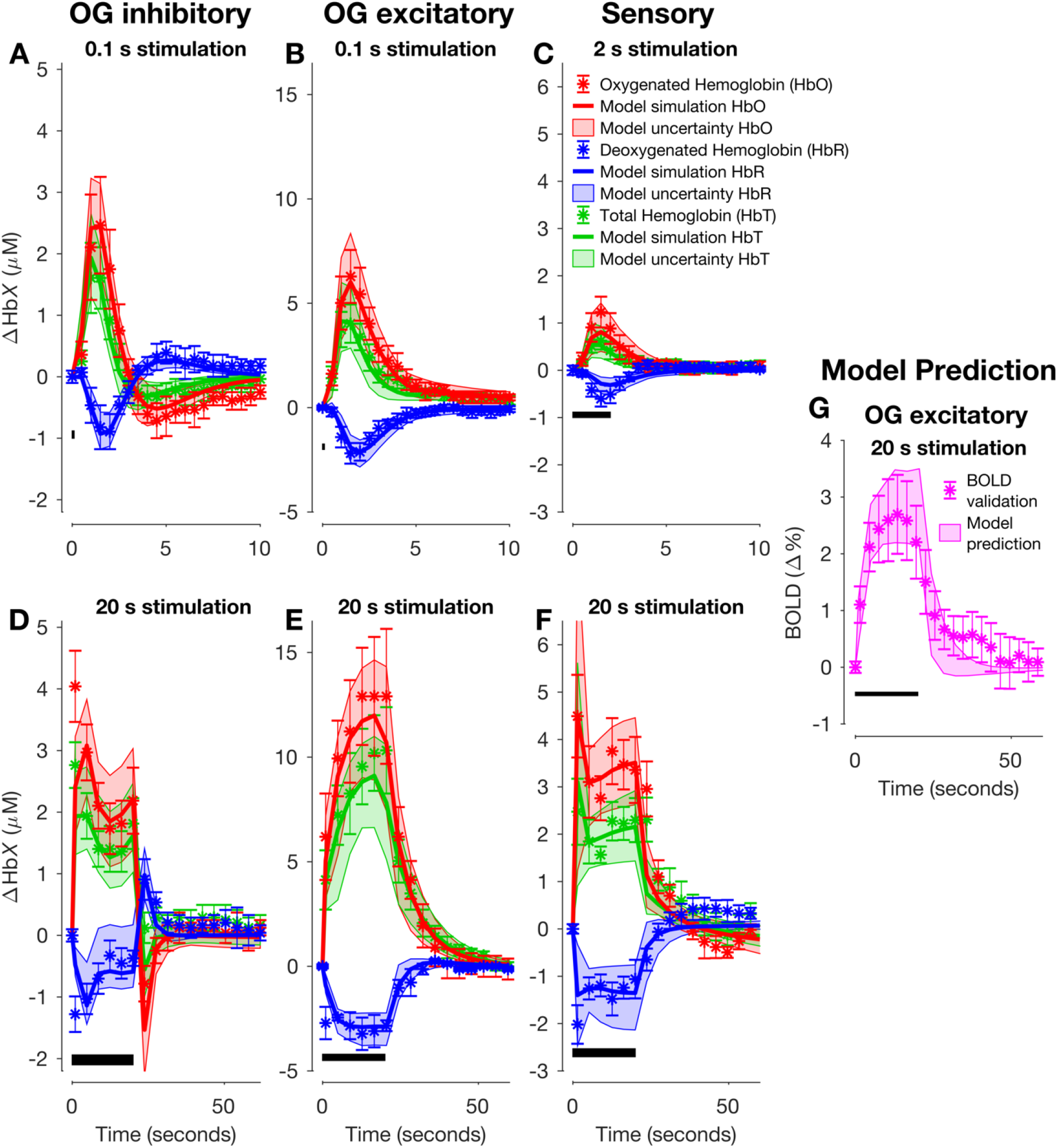
Model estimation to experimental data of hemoglobin changes in mice for three different stimulation types: optogenetic (OG) activation of inhibitory (**A & D**) and excitatory (**B & E**) neurons, and a sensory stimulation (**C & F**). For each stimuli type, a short stimulus (OG: 100 ms of light (**A−B**); sensory: 2 s (**C**)) and long stimulus (OG: 20 s (**D−E**); sensory: 20 s (**F**)) was used. This is denoted with the black bar in the bottom left portion of each graph. The shown experimental data are replotted versions of Fig. S5 in the study by Desjardins *et al*. [76]. For each graph: experimental data consisting of oxygenated hemoglobin (HbO; red symbols), deoxygenated hemoglobin (HbR; blue symbols), and total Hb (HbT; green symbols); the uncertainty of the experimental data is presented as SEM (colored error bars); the best model simulation is seen as colored solid lines corresponding to respective measurement variable; the model uncertainty as colored semi-transparent overlays; the x-axis represents time in seconds, and the y-axis is the change in hemoglobin concentration (μM). **G**: Model predictions (shaded area) and experimental data (mean ± SEM, symbols) of a BOLD response to an identical stimulus is shown in **E**. The validation experimental data were extracted from Desjardins *et al*. 2019 [76].

**Figure 6.**
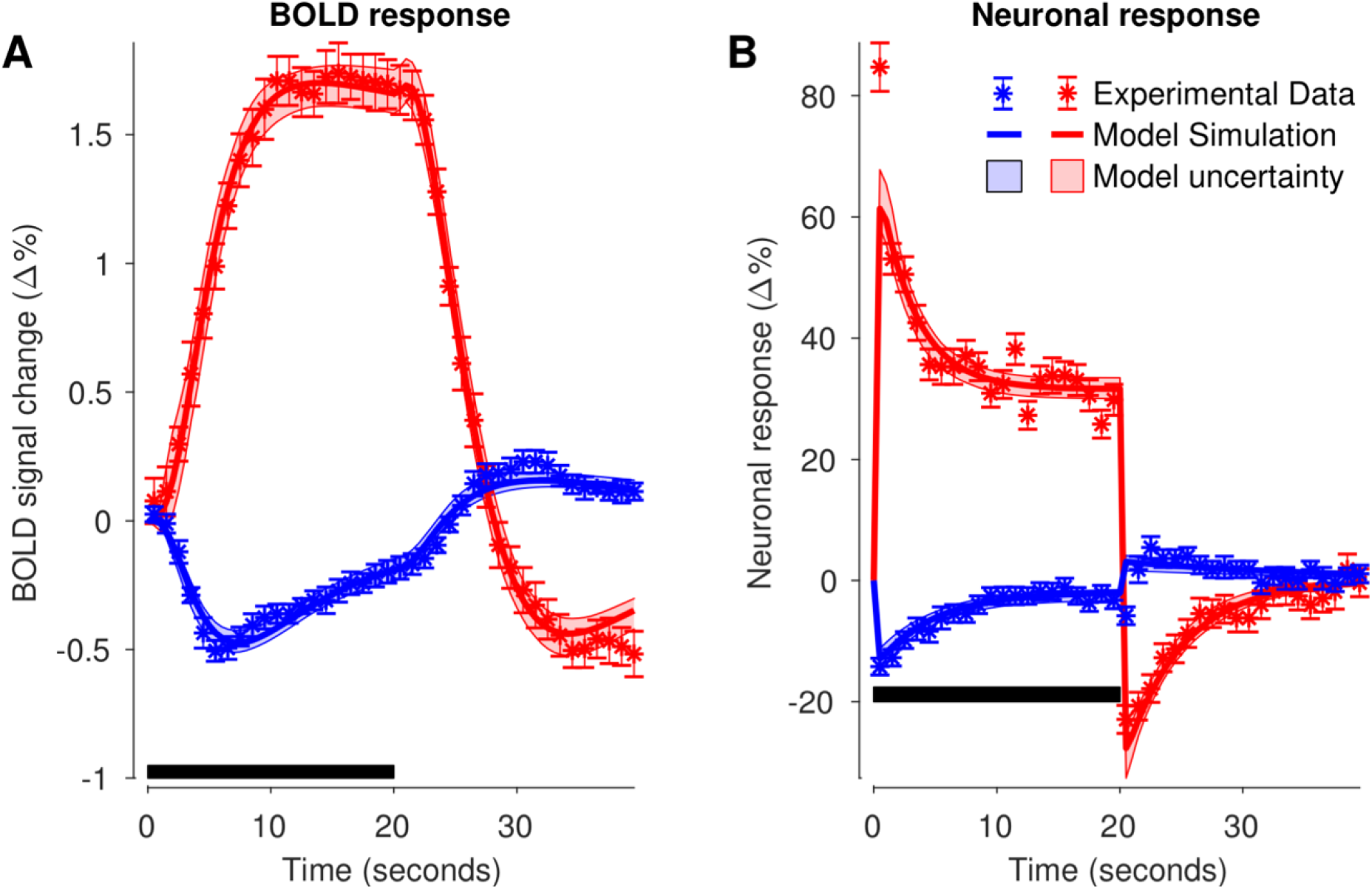
Model estimation to experimental data of BOLD (A) and electrophysiological (B) responses in macaques for two variants of a visual task (blue and red symbols). The task was a 20 s visual stimulation of V1 in the visual cortex (marked with the black bar in the bottom left portion of each graph). For each graph: experimental data (colored symbols); The uncertainty of the experimental data is presented as SEM (colored error bars); the best model simulation seen as colored solid lines corresponding to respective measurement observables. The model uncertainty is seen as the semi-transparent colored areas. The x-axis represents time in seconds, and the y-axis is BOLD signal change (Δ%) for **A** and LFP expressed as percent change from baseline (Δ%) for **B**. The experimental data were extracted from Fig. 1D & 2A from Shmuel *et al*. 2006 [10].

### 2.5 Model implementation

Implementation and simulation of the model equations (specified in Section 2.2) were carried out using the ‘Advanced Multi-language Interface to CVODES and IDAS’ (AMICI) toolbox [79, 80]. Parameter estimation for the model evaluation was carried out using the MATLAB implemented ‘MEtaheuristics for systems biology and bioinformatics Global Optimization’ (MEIGO) [81] toolbox and the enhanced scatter search (eSS) algorithm. In addition, eSS was used in conjunction with two local optimization solvers: dynamic hill climbing [82] and the interior point algorithm included in FMINCON (MATLAB and Optimization Toolbox Release 2017b, The MathWorks, inc., Natick, Massachusetts, United States) paired with the calculation of the objective function gradient using AMICI. For the model uncertainty analysis, we employed the ‘Parameter EStimation ToolBox’ (PESTO) [83] and the region-based adaptive parallel tempering algorithm [84] to generate the posterior distributions of the parameters. The parameter bounds were [-4.5,4.5] in log_10_ space unless specified otherwise in Section 2.2.

### 2.6 Data and code availability

The code necessary for reproducing all the figures and results reported herein is provided in the Supplementary material as MATLAB scripts.

## 3 Results

Below, experimental data gathered from five studies: Drew *et al*. [75], Desjardins *et al*. [76], Uhlirova *et al*. [55] (Section 3.1), Shmuel *et al*. [10] (Section 3.2), and finally, Huber *et al*. [77] (Section 3.3) are presented with corresponding model simulations including the uncertainty of the model.

### 3.1 NVC measures in awake and anesthetized mice

#### 3.1.1 Fractional diameter change of arterioles and venules in awake mice

The model was trained using experimental data extracted from the study of Drew *et al*. [75], consisting of arteriolar (Fig. 3A–C, red symbols) and venular (Fig. 3D–F, blue symbols) diameter changes induced by a sensory whisker stimulation for three different durations: 20 ms (Fig. 3A, D), 10 s (Fig. 3B, E) and 30 s (Fig. 3C, F). The stimulation consisted of air puffs (8 Hz, 20 ms duration), and just as in Barrett *et al*. [34], we opted to implement this paradigm as one block with the following durations: 125 ms, 10 s, and 30 s. In the model, we calculate the diameter change as the square root of the volume change for each respective compartment. The model parameters were fitted to all experimental time series simultaneously (Fig. 3A–F, colored symbols), achieving a quantitative acceptable agreement to experimental data (Fig. 3A–F, solid colored lines), evaluated using a *χ*^2^-test (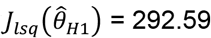, cut-off: *χ*^2^ (288 data points) = 328.58 for *α* = 0.05). The degrees of freedom for the confidence interval was *df*_*H*1=_38, which is equal to the number of estimated parameters. The estimated model uncertainty is depicted in the form of shaded red and blue areas in Figure 3.

The simulations follow the qualitative behavior of the experimental data for each graph except for Fig. 3E, which is the venular response to a 10 s stimulation. For that stimulation, the model simulations show a slower rise and decline than the experimental data (Fig. 3E, compare blue line with blue symbols). A more accurate fit can be obtained by allowing the viscoelasticity and stiffness coefficients of the capillary and venous compartments to change between the two short (125 ms & 10 s) and the long (30 s) stimulation (Fig. S1, 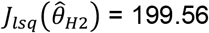). This improved agreement indicates that a non-linear vascular model is needed to fully capture the dynamics seen in this data set. For the other experimental time series, the model simulations closely resemble the experimental data, and the model uncertainty enwraps most of the experimental data points.

#### 3.1.2 The dynamic behavior observed can be explained by two distinct dilation processes

The arteriolar responses transition from a one peak (Fig. 3A) to a two-peak response (Fig. 3C), as the stimulation length increases. In other words, for the short stimulation, the response rises to a peak amplitude value around 2–3 seconds, followed by a decline to baseline. For the two longer (10 s and 30 s) stimulations, a quick initial dilation is followed by a small cessation to a plateau value around 10 s (Fig. 3B–C). At 10 s, the stimulation ends in one case, with a minor dilation before returning to baseline diameter (Fig. 3B). However, for the 30 s stimulation, a new dilation phase occurs, consistently dilating until the end of the stimulation (Fig. 3C, t = 10–30 s). Finally, after stimulus cessation for the longest stimulation, but not for the shorter stimulations, a post-peak undershoot in the arteriolar diameter is observed before returning to baseline (Fig. 3C, t ∼ 50 s).

As seen in Fig. 3A–C, these complex dynamic behaviors between different stimulation lengths are captured by the model. It is therefore interesting to examine by which mechanisms the model produces these behaviors. In Figure 4, we illustrate how the model simulations can provide such an examination. Model simulations are shown for all three stimulation lengths: 0.125 ms (Fig. 4A & E), 10 s (Fig. 4B & F) and 30 s (Fig. 4C & G). Furthermore, the top three figures correspond to the rapid neuronal responses (Fig. 4A–C), and the bottom three figures correspond to the impact of the vasoactive substances: NO, PGE_2_, NPY (Fig. 4E–G). In addition, the simulations are shown for all three neuronal types and corresponding vasoactive substances: pyramidal neurons and PGE_2_ (green), NO interneuron and NO (blue), and NPY interneurons and NPY (red). Finally, the resulting behavior (Fig. 4E–G, black dashed line) is produced by the summation of the three impacts, where NPY is vasoconstrictive (negative), and PGE_2_ and NO are vasodilative (positive). To help navigate these different simulations, we have placed a simplified view of the model in the middle (Fig. 4D).

With these interpretations in place, we can now understand how the model produces the complex dynamic behaviors observed in data. It is easiest to follow the chain of events by starting at the last step before the observed behavior. The response to a short stimulation is produced by the fast release of NO, which produces the initial peak in the arteriolar response (Fig. 4F–G, blue line) and which thereafter simply declines. The other two signaling pathways are slower and therefore negligible. Also for the longer stimulations, the NO behavior is important. For these stimulations, the NO response culminates quickly, where after the impact of NO falls which causes a decrease after the first peak.

However, for longer stimulations this decrease is replaced by a new increase, which is caused by the rise of the slower PGE_2_ signaling arm from the pyramidal neuron (Fig. 4F–G, green line). In other words, the PGE_2_ arm is responsible for the second dilation phase observed for the 30 s stimulation (Fig. 4G, green line). Finally, the NPY signaling arm is only important to explain the post-stimulus undershoot observed in the 30 s stimulus paradigm as it is so slow that it requires such long stimulation to become larger than the two other arms (Fig. 4G, red line is above the blue and green for t ∼ 50 s). In summary, the initial peak is explained by the fast NO arm, the second peak is explained by the slower PGE_2_ arm, and the post-peak undershoot is explained by the even slower NPY arm.

These different dynamics in the three arms occur at the downstream secretion level, but must be initiated by corresponding changes at the rapid neuronal level. This rapid level is depicted in Figure 4A–C. For the shortest stimulation (Fig. 4A), the NO interneurons are here the fastest, and pyramidal neurons and NPY interneurons are equally fast. For the two longer stimulations (Fig. 4B–C), the electrophysiology reaches a steady-state within 1 second, where-after no changes occur until the stimulation ends. Since the secretion of NPY is much slower than PGE_2_, and since both these dynamics occur over many seconds, the differences in dynamics at the downstream secretion level is almost exclusive explained by the difference in intracellular signaling. The only difference concerns the NO dynamic, which has a peak and decline at the secretion level, which is caused by a corresponding rise and fall at the electrophysiology level.

#### 3.1.3 The model can describe and correctly predict arteriolar responses from Uhlirova et al. [55]

In our previous computational work (see [41]), we developed a model capable of estimating and predicting the experimental data found in Uhlirova *et al*. [55]. To verify that this extended model still maintains a good agreement and predictive power to that experimental data, we re-did the model training and model predictions in [41]. The Uhlirova study reports arteriolar diameter changes in mice evoked by different stimuli. These different stimuli include for instance OG and sensory stimulation, where OG can stimulate either pyramidal neurons or inhibitory interneurons separately. They do this cell-specific stimulation with and without pharmacological perturbations, using the NPY receptor inhibitor BIBP, and the glutamatergic signaling inhibitors CNQX and AP5. These pharmacological perturbations cut off the cross-talk between NPY and the vasculature and the cross-talk between pyramidal neurons and interneurons, respectively. Furthermore, they also study the animals during awake and anesthesia conditions. In [41], our model can explain all this data with the same parameters, and can also predict independent data not used for training. Our updated model has the same capability (see Supplementary Material S1.2; Fig. S2), and these results *e*.*g*., support the notion that NPY acts vasoconstrictively, and causes the post-peak undershoot.

#### 3.1.4 Hemoglobin and BOLD measures in awake mice

Next, we analyzed experimental data from the study of Desjardins *et al*. [76]. In brief, the study reports hemodynamic measures of blood oxygenation (HbO, HbR, HbT) from three different stimulations: 1) OG stimulation of inhibitory interneurons; 2) OG stimulation of pyramidal neurons, and 3) sensory whisker stimulation using air puffs. Each of these three paradigms had a short (OG: 100 ms light pulse; sensory: 2 s of air puffs at 3–5 Hz) and long (OG: 20 s of 100 ms light pulses at 1 Hz; sensory: 20 s of air puffs at 3–5 Hz) duration. They also reported fMRI-BOLD data for the long OG stimulation of pyramidal neurons. We implemented these stimulus paradigms in the model and assumed a 100 ms air puff delivered at 3 Hz for the sensory stimulus. To capture the variability in frequency, which gives rise to an observable difference in stimulation strength between the two sensory experiments (Fig. 5C, F), we allowed the input parameters (*k*_u,i_) to change between the short and long duration of the sensory stimulation but could keep the same values for both of the OG paradigms (see Discussion 4.2). Furthermore, in the model, we assumed a baseline blood volume concentration of 100 μM hemoglobin, in line with previous models [35, 37, 85].

With these data and model settings, the model was trained to experimental data of hemoglobin dynamics for all three stimulation paradigms simultaneously (Fig. 5A–F, red/blue/green symbols), and the BOLD data was left out to be used for model validation (Fig. 5G, magenta symbols). The model achieved a quantitative acceptable agreement to experimental data (Fig. 5A–F, red/blue/green lines) after parameter estimation (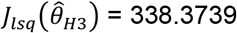, cut-off: *χ*^2^ (354 data points) = 397.87. The degrees of freedom for the confidence interval was *df*_*H*4=_43, which is equal to the number of estimated parameters (Fig. 5A–F, shaded areas).

To validate the model and avoid overfitting of the parameters, we used the model to generate predictions of a BOLD response for a 20 s OG stimulation of pyramidal neurons. These predictions are depicted as the magenta shaded area in Fig. 5G. The corresponding experimental data (Fig. 5G, magenta symbols) lie within the model prediction bounds. The validation of the model is supported by a *χ*^2^ test, which does not reject the model with respect to validation data (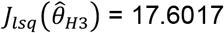, cut-off: In summary, all results up until now taken *χ*^2^ (23 data points) = 35.1725).

In summary, all results up until now taken together show that our model structure provides an acceptable explanation for a wide variety of aspects observable in mice. Figures 3–4 unravel the role of three regulatory arms: NO interneurons are responsible for the rapid rise, PGE_2_ from pyramidal neurons are responsible for the second peak that occurs for long stimulations, and NPY interneurons for the post-peak undershoot. All these results are using vessel diameter as a proxy for the NVC control. In Figure 5, we confirm that the same model can explain the NVC control seen using other measures: total hemoglobin serves as a proxy for blood volume, HbO/HbR examines the balance between metabolism and blood flow, and the BOLD signal provides a link to the most common non-invasive measure in primates and humans.

### 3.2 Neuronal and BOLD measures in anesthetized macaques

#### 3.2.1 The model agrees with BOLD and LFP data

We analyzed experimental data from the study of Shmuel *et al*. 2006 [10], which presents unique data for higher primates, namely macaques. We extracted electrophysiological and BOLD responses induced by a visual task in anesthetized macaques. The task was presented in two ways, inducing both positive and negative responses in the same area (see Section 2.4.3 for details). These data were extracted from Fig. 1D and 2A from the original manuscript. The electrophysiological response was related to the local field potential (LFP), which is to a large extent, influenced by pyramidal neuron firing (see review in [86]). Therefore, we assumed the following measurement equation for the model:

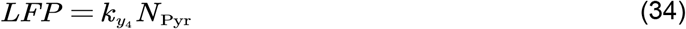

where N_Pyr_ is the phenological activity state of pyramidal neurons, and 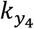 is an unknown scaling constant. Between a positive and negative response, we allowed the following parameters to be changed: the input stimulus parameters (*k*_u,Pyr_, *k*_u,NO_, *k*_u,NPY_), where *k*_u,Pyr_ 0, to be able to have both positive and negative values; and the parameters governing the inter-neuronal interactions between N_Pyr_, N_NO_, and N_NPY_ (*k*_PFY_, *k*_PF2_, *k*_INF1_, *k*_INF2_, *k*_IN1_, *k*_IN2_).

The model was trained to experimental data for both positive (Fig. 6A–B, red symbols) and negative (Fig. 6A–B, blue symbols) responses simultaneously. The model achieved a quantitative acceptable agreement to experimental data (Fig. 6A–B, red and blue solid lines) after parameter estimation (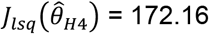, cut-off: *χ* ^2^ (160 data points) = 190.52). The degrees of freedom for the confidence interval was *df*_*H*4=_49, which is equal to the number of estimated parameters (Fig. 6A–B, shaded areas).

As seen, the simulations agree overall well with the experimental data, as almost all experimental data points lie within the model uncertainty area (Fig. 6A–B, compare shaded areas with corresponding symbols). However, for the positive response, the model fails to agree with the first data point of the LFP experimental data during stimulation, not reaching the same amplitude (Fig. 6B, red, t = 1 s). This issue can be mitigated by allowing the parameters governing the neuronal states N_Pyr_, N_NO_, and N_NPY_, to change between stimulus and post-stimulus periods (as previously done in the computational work of [36], see Discussion 4.2).

### 3.3 MR-based hemodynamic measures in awake human

#### 3.3.1 The model can describe data used for training and predict independent validation data

Further, we looked at the study conducted by Huber *et al*. [77]. We extracted the experimental measurements of CBV, CBF, and BOLD responses in humans, evoked by multiple visual stimuli. The study employed a set of three different flickering checkerboard patterns as the visual stimuli (see Section 2.4.5. for more details). The extracted data originated from figures 4A−B, 5B−C, and 6C in the original paper [77]. The model was trained on experimental data consisting of both positive and negative CBV (Fig. 7A) and BOLD responses (Fig. 7D), as well as experimental data for the total CBV changes for the excitatory (Fig. 7B, black) and inhibitory (Fig. 7C, black) tasks. As in Section 3.2, we allow the input stimulus parameters (*k*_u,i_) to change between the three tasks, and the parameters governing the inter-neuronal interactions (*k*_PF1_, *k*_PF2_, *k*_INF1_, *k*_INF2_, *k*_IN1_, *k*_IN2_) to change between positive and negative responses. Parameter estimation was employed to achieve an acceptable fit to the data (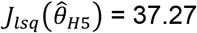, cut-off: *χ* ^2^ (122 data points) = 148.78). The simulation uncertainty can be seen as the colored shaded areas in Fig. 7A–E. As seen, the model simulations have a good agreement with the experimental data (Fig. 7A and D, shaded areas, and Fig. 7B–C, black shaded areas), as the model uncertainty area overlaps well with the experimental data points. To test the model’s predictive qualities, we used the model to generate predictions of CBF for both the positive and negative responses (Fig. 7E), as well as predictions of arterial CBV and venous CBV for both the excitatory (Fig. 7B, red and blue) and inhibitory flickering checkerboard tasks (Fig. 7C, red and blue). There is a good agreement between the model predictions and the experimental data.

**Figure 7.**
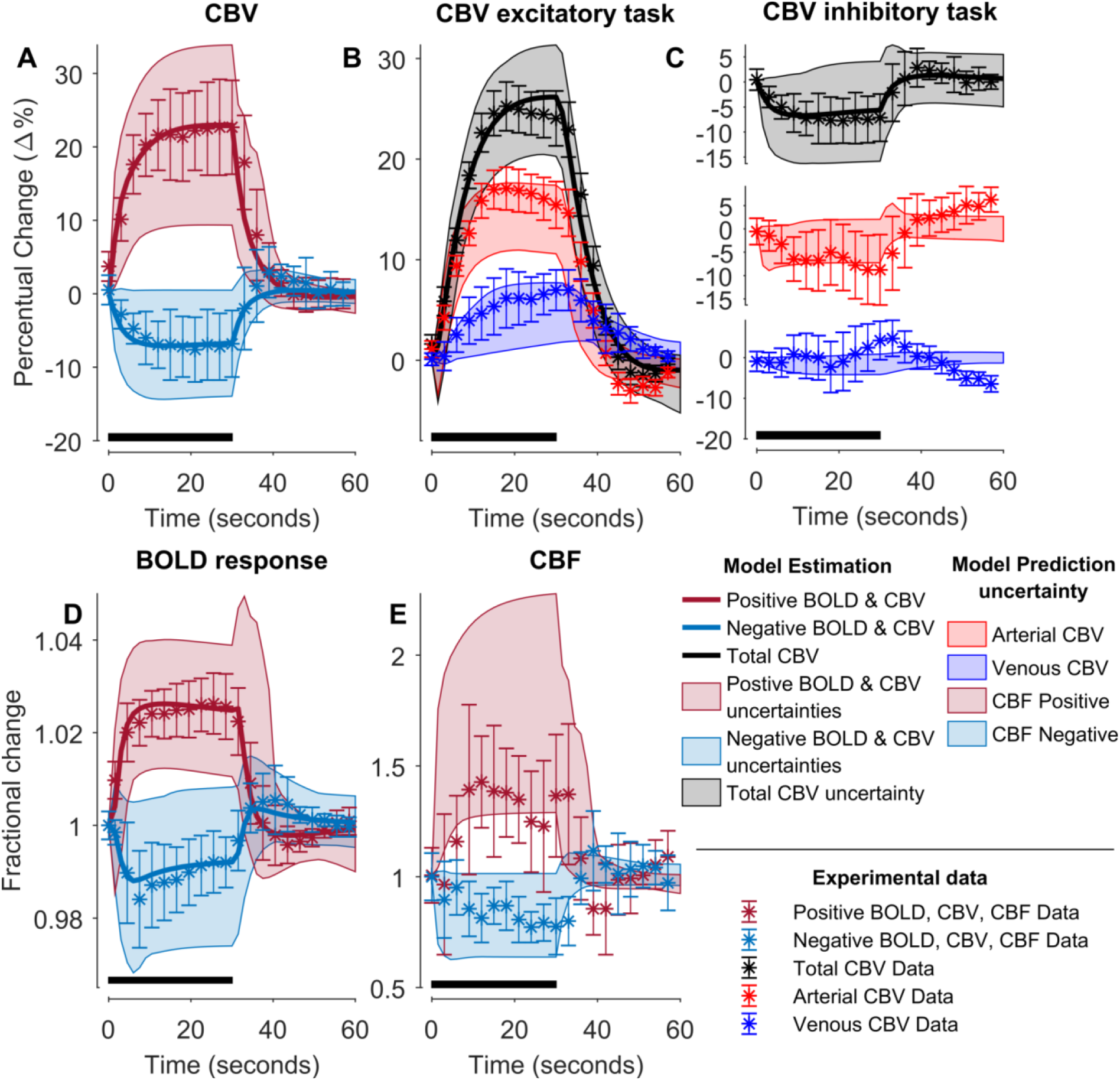
Model estimation and prediction to MR-based experimental data in humans for different visual stimulation tasks: The graph contains data for three different tasks: CBV (**A**), BOLD signal (**D**), and CBF (**E**) changes for small flickering checkerboard task, and compartment-specific CBV changes for excitatory (**B**) and inhibitory (**C**) flickering tasks. For each graph, the stimulus duration (30 s) is depicted the black bar in the bottom left portion of each graph; experimental data shown with the measurement uncertainty as SEM (colored error bars); the best model simulations is seen as colored solid lines corresponding to respective measurement observables; the model uncertainty is depicted as colored semi-transparent overlays; the x-axis represents time in seconds, and the y-axis is the change in fractional (**D, E**) or percentual change (**A–C**) of the measurement observables. The model prediction uncertainty of CBF (**E**) and compartment-specific CBV changes (**B, C**) is depicted as semi-transparent overlays. The experimental time series are taken from Huber *et al*. [77].

## 4 Discussion

Currently, no existing model of the NVC can simultaneously describe most available NVC data (Fig. 1B, 1D): 1) BOLD, 2) CBV, 3) CBF, 4) Hemoglobin measures, 5) optogenetic and sensory stimulations, 6) pharmacological perturbations including the effect of anesthetics, and 7) phenomenological description of LFP-related signals. Herein, we provide a first such model (Fig. 2). The model can quantitatively explain experimental data used for model training (Figs. 3, 5, 6, 7, S2A– E) and correctly predict experimental data not used for training (Fig. 5G; Fig. 7BCE; Fig. S2F–H), from rodents (Figs. 3, 5, S2), primates (Fig. 6), and humans (Fig. 7). This validated model provides a first interactive and inter-connected explanation for how the complicated dynamics and interdependencies in all of these data can be understood. We illustrate this potential by using the model to explain the double peak in arteriolar diameter in response to prolonged stimuli (Fig. 4G, black broken line): the first peak is produced by the rapid NO dilating arm (Fig. 4E–G, blue line), the second peak is produced by the slower PGE_2_ arm (Fig. 4E–G, green line), and the post-peak undershoot is explained by the even slower NPY arm (Fig. 4F, red line). This new quantitative understanding can also serve as a foundation for a new more integrative approach to analysis of neuroimaging data, which opens the door to many new applications in both basic science and in the clinic.

### 4.1 Models of the NVC

The NVC has previously been mathematically modeled using many different approaches. The earliest models, which still are used in the conventional analysis of BOLD-fMRI data today, are based on the general linear model (GLM), as implemented in software packages like statistical parametric mapping (SPM) [87, 88]. GLM allows one to identify brain areas with activity, but does so in a way that does not make use of any biological understanding, and thus cannot incorporate any additional data. Such incorporations required more mechanistic models. Simple mechanistic models were presented by Friston *et al*. [27, 89] and Buxton *et al*. [28, 29]. These models are commonly referred to as the Balloon model. The Balloon model has become widely used, spawning many subsequent variants and improvements [36, 90–93]. These models typically only describe venous volume changes (as this is the dominant contributor to the BOLD-fMRI signal). One significant shortcoming with these models is that they do not consider the arteriolar volume that is actively regulated in the NVC, and that it primarily is the arteriolar volume that changes upon short event-related stimuli (Fig. 3). To remedy this, some have developed Windkessel-like models, which are electrical circuit equivalent models of pressures, volumes, and flows [34, 35, 94] that can describe all cerebrovascular volumes. In contrast, others have instead opted to exclusively describe arteriolar volume changes, which only are applicable for short stimuli [95, 96]. Finally, other types of models are based on partial differential equations and detailed diffusion images [97–100]. A common shortcoming for all these mentioned approaches is that these models lack biochemical mechanisms underpinning the intracellular signaling, which translates neural activity to hemodynamic changes. Mathematical models describing these intracellular processes found in the NVC exist [38–40, 101–104], but they are typically connected to the more simplified Balloon model, or lack realistic descriptions of the vascular dynamics. In conclusion, no model that is both based on biochemical mechanisms and which propagates vascular changes using a Windkessel model has previously been described (Fig. 1D). In this work, we present a first such model.

### 4.2 Discussion on problematic data points

While the overall agreement between model and data is statistically acceptable, as the model simulations pass *χ*^2^-tests, there are some individual data points that cannot be explained by the model and that warrant some additional comments. In Section 3.1.1, we present the model estimation to experimental data collected from the original research in [75]. While the overall model agreement overlaps well with the experimental data, the model simulation does not agree well with data of venule dilation for the 10 s stimulus (Fig. 3E, compare symbols with the shaded area). The experimental data exhibits a faster post-stimulus decay, while the venule dilation persists for a longer period of time in the model simulation. The identical behavior is seen in the original manuscript for the vascular model [34] (see Fig. 2F in [34] for the corresponding simulation), which was compared to the same experimental data. The reason why both these models fail to accurately capture the dynamics in Fig. 3E is that the 10 s stimulus data show a much faster decline after the peak than the 30 s stimulus data. In the original paper for the vascular model, Barrett and co-workers speculate that this discrepancy in data is due to few measurements on single vessels, and that the fast signal decay in 10 s stimulus data would disappear if averaged over many vessels. Another confounding factor reported along with the experimental data ([75]) is that changes in venule diameter of less than 2 %, and capillary diameter changes of less than 7 %, could not be captured using their measurement setup. For this reason, the data in Fig. 3D–E are at or below the detection limit, and we therefore view the disagreements with corresponding model simulations to not be a major concern.

In Section 3.1.4, the model simulation fails to reach the first data point in Fig. 5D (HbO, HbR, HbT) and Fig. 5E (HbR). This failure is due to the following inconsistency in the data. Examining the difference between the short and long OG IN stimulation for HbO, we would assume that the first data points up until t = 1 s would be approximately identical, as only one light pulse should have been emitted in both paradigms. However, for t =1 s, experimental data points differ by a factor of two. As we assume that the effect of each light pulse is identical between short and long stimulation (since the model does not know the future), the model cannot describe this behavior. For this reason, we choose to keep the model with this shortcoming, since the alternative would be to have different parameters for short and long OG stimulation.

In Section 3.2.1, the model describes all data points except for the first data point in the LFP data for the positive response (Fig. 6B, red). In previous computational work based on the same data [36], parameters were allowed to change between stimulus and post-stimulus periods. This alternative approach gives a satisfactory agreement with data, including the first data point. However, we see no reason for why the parameters should change when the stimulus ends, and we therefore opt to accept the disagreement with the first data point. It is possible that a more non-linear description of LFP would provide a better agreement with this first data point. Again, we choose not to follow such a path, as it might lead to overfitting. A similar disagreement with experimental data can also be observed for the negative LFP response (Fig. 6B, blue). Here, the model simulations instantly return towards baseline upon stimulus cessation, while the experimental data show a ∼ 1 s delay before rapidly changing. In Havlicek *et al*. [36], this was compensated for by simply prolonging the stimulation period by 1 s. If our model allowed for a similar prolonging, the same improvement would also be seen in our simulations.

In Section 3.3.1, the model prediction uncertainty for the positive response of CBF has a high uncertainty, and therefore some parts of the prediction uncertainty lies outside of the data (Fig. 7E, red). Further, there might be dynamics in the changes of venous CBV for the inhibitory task, not captured by the model. However, because of the high uncertainty of the data, this discrepancy cannot be concluded (Fig. 7C, blue).

### 4.3 Limitations with the current model

As with all models, there exists a set of limitations and assumptions which are necessary, but still limit the scope of conclusions that can be drawn using the model.

First, our electrical activity description is simplified. More specifically, we use a simple phenomenological relationship between GABAergic interneurons and pyramidal neurons, where pyramidal neurons excite the GABAergic interneurons, which in turn act to inhibit the pyramidal neurons (Eq. 3). An important development would be to integrate this model with common simulators of electrical activity, such as NEURON [105] or NEST [106] (see review in [107]). These models describe how membrane potential fluctuations arise as the result of the opening and closing of ion channels due to glutamatergic and GABAergic signaling. Such an integration would also allow for the description of individual neurons, as there exist structurally and functionally different morphologies of interneurons (see [108] for review), which would be a more realistic description compared to our current lumped approach.

Second, since we describe a wide variety of data, our choice of using the same model structure for all the data has implied some choices. For each experimental data set used in this study, we fitted some of the model parameters in the model specifically to each experimental study, but kept all parameters the same within the same study. More specifically, all parameters are the same for all simulations in Figs. 3–4, 5, 6, 7, S1–S2, but some parameters are different, for instance between Figs. 3 & 5. Details on this is given in supplementary model files. Some other parameters are not fitted to any of these data but are assumed to be the same across studies. For instance, the vascular volume fractions V_*i*,0_ and baseline oxygen transport parameters (see Section 2.2.4) are assumed to be the same across the different species. Furthermore, some experimental modalities measure directly on individual arterioles and/or venules (Figs. 3 & S2), while other modalities measure a lumped estimate over the entire arterial or venous compartment (Figs. 5–7). In the model, we use the same simulation variable to describe both individual and lumped measurements.

Another highly simplified component of our model concerns metabolism. While the metabolism of oxygen occurs in the model, with both a baseline and a stimulus-dependent component, this part could, in principle, be expanded to incorporate the commonly measured metabolites in glia. These metabolites are typically measured using magnetic resonance spectroscopy (MRS), and models capable of describing these interplays already exist [109, 110]. In addition, there are some known additional mechanisms of the NVC that we have not included. For instance, we do not include the role of astrocytes. Astrocytes have end-feet ideally placed around smooth muscle cells on arterioles and pericytes, and astrocytes are known to regulate the basal tone of arteries [111–114]. The reason we did not include these is that the exact role is unclear, and we did not find useful dynamic data. Another vasoactive pathway involves inward-rectifier K^+^ (K_IR_2.1) channels located on capillary endothelial cells, which open upon extracellular increases of K^+^ as a result of neural activity [115]. The opening of these channels induces hyperpolarization, which propagates rapidly upstream to arteries [115]. We do not include direct actions on the capillaries, as reported in [50, 116].

Lastly, in the model, the amount of some states (Eq. 3-7) are specified in arbitrary units, which means that we describe biological values such as concentrations, scaled by unknown scaling constants. Furthermore, the vascular states (Eq. 9–11) are normalized to a baseline value of 1, in order to improve computational stability and simplify calculations. This particular simplification is inherited from Barrett *et al*. [34, 42, 43]. Despite these renormalizations, our model can still be used to predict and simulate observations that are possible to capture in experimental data.

### 4.4 Strengths and potentials

The big potential with our model is not only the incorporation of more experimental data than any previous model (Fig. 1D), but that we bypass uncertain parameter values and thus can provide predictions with uncertainty, which can be useful in a variety of contexts. One of the main advantages of these predictions is that they can be used to understand mechanisms, which is illustrated in Figures The simulations in Figure 4 are given by a single parameter vector, producing a single line. This is appropriate because the only question we ask is how the complex bimodal response can be produced by the model. For other types of questions, we want to plot predictions with uncertainty.

One often seeks well-determined predictions, sometimes called core predictions [104, 117], and they are useful for instance when validating the model and designing new experiments [118]. Such validations are done for instance in Fig. 5G or Fig. 7BCE (areas without solid lines). These core predictions can also be used to measure otherwise non-measurable entities, which can be estimated from a combination of available data. One classical example of such model-based “measurements” is the approximate measurement of metabolism, which can be done using the Davis model [119], which uses the combination of BOLD and CBF data. In our much more comprehensive and complete model, many more different data sets can be combined, and more possible features can be estimated. These estimated features can both aid in the understanding of mechanisms in basic research (Fig. 4) and be used to identify new biomarkers which can be used to stratify between different patient groups in the clinic.

## 5. Author contributions

SS, FE, ME and GC designed and conceived the study. SS carried out the model formulation, data extraction, numerical simulations and analysis, and wrote the draft. HP carried out data extraction and model formulation under supervision of SS. NS performed numerical simulations and analysis under supervision of SS. HP, NS, FE, ME, and GC provided critical revisions and comments on the manuscript.

## Supplementary material accompanying

The supplementary material contains the following: additional model estimations, and the posterior probability profiles of the parameters for all model estimation.

### S1 Additional model estimations

#### S1.1 Allowing the stiffness and viscoelasticity parameters between the capillary and venular compartments to change between 10 s and 30 s stimulation for data from Drew et al. [1]

**Figure S1.**
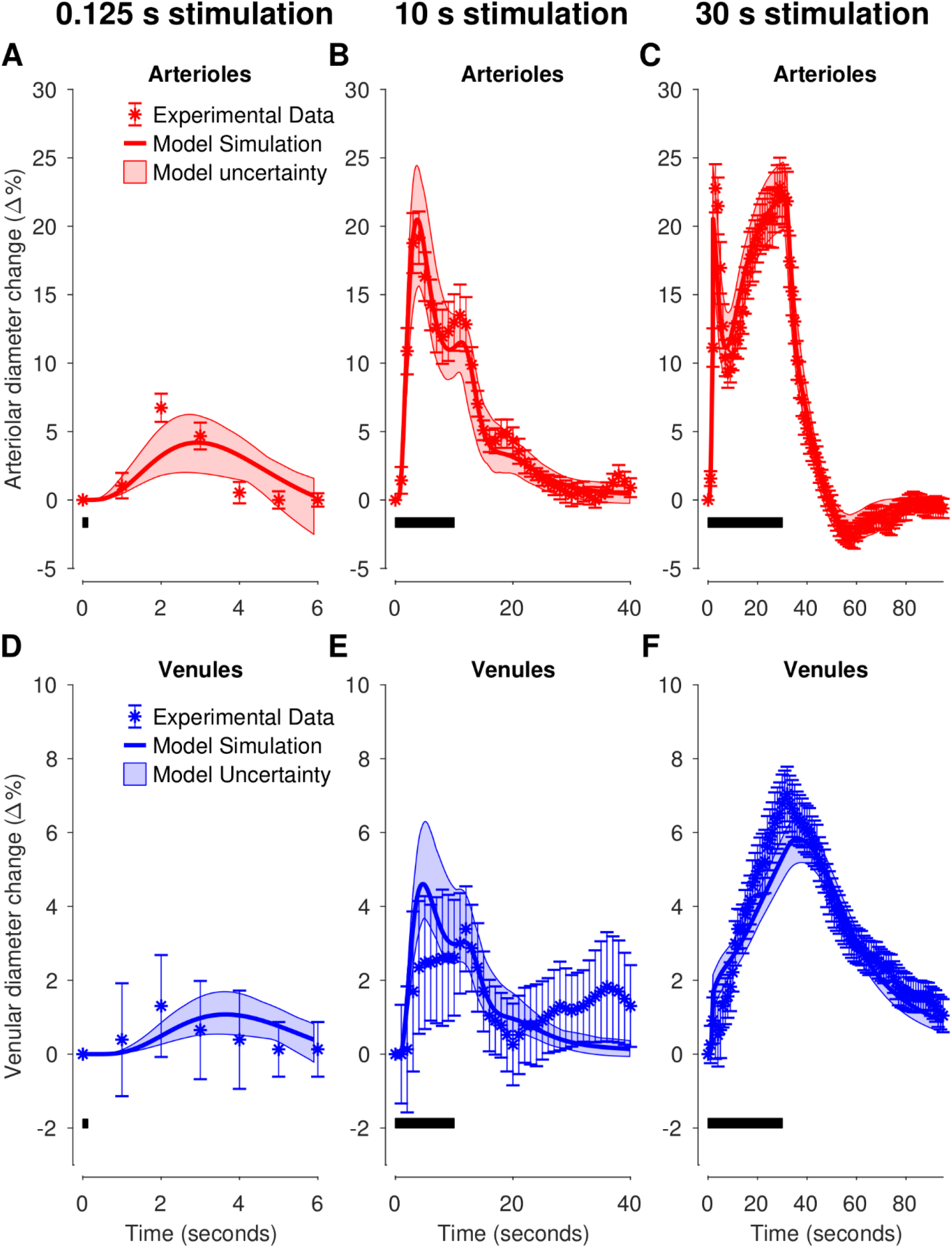
Model estimation to experimental data of arteriolar (**A–C**) and venular (**D–F)** volume changes in awake mice for three different sensory stimulation lengths: 125 ms (**A & D**), 10 s (**B & E**), and 30 s (**C & F**). The viscoelasticity and stiffness coefficients of the capillary and venular compartment change at t = 3s for the long stimulation (**C & F**). Experimental data are replotted versions of data presented in Fig. 2C of the original manuscript by Drew *et al*. [1]. The stimulation lengths are denoted with the black bar in the bottom left portion of each graph. For each graph: experimental data (colored symbols); The uncertainty of the experimental data is presented as SEM (colored error bars); the best model simulation is seen as a colored solid line; the model uncertainty as colored semi-transparent overlay. The x-axis represents time in seconds, and the y-axis is the normalized vessel diameter change (Δ%).

#### S1.2 Model estimations for data from Uhlirova et al

The Uhlirova study (see [2]) reports arteriolar diameter changes in mice evoked by different stimuli for awake and anesthesia conditions. To account for the anesthetic effect, we allowed 12 parameters governing the neuronal interactions to change between awake and under anesthesia: the three stimulus-scaling parameters *k*_u,i_; the three decay rate parameters *sink*_N,i_ affecting N_i_, and finally the six neuronal interaction parameters *k*_PF1_, *k*_PF2_, *k*_IN1_, *k*_IN2_, *k*_INF1_, and *k*_INF2_. Furthermore, we allow the stimulus scaling parameters *k*_u,i_ to change between OG and sensory stimulation. See our previous study for more details [3].

The thusly parameterized model was trained on experimental data consisting of all control conditions from the Uhlirova *et al*. [2] (Fig. S2 left panel A–E, black symbols). The model parameters were fitted to the experimental data, achieving an acceptable agreement with data (Fig. S2 A–E solid thick red lines), evaluated using a *χ*^2^-test (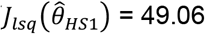 cut-off: *χ*^2^ (190 data points) = 223.16 for *α* = 0.05). Next, the uncertainty of the model was approximated using Markov chain Monte Carlo sampling (10^5^ iterations), producing posterior parameter profiles (Fig. S5) and a sample of all *χ*^2^ acceptable parameters. A confidence interval (Eq. 33 in the main manuscript) with *α*=0.05 and degrees of freedom *df*_*H*3=_54 (equal to the number of estimated parameters) was drawn and can be seen as the red shaded areas in Fig. S2A–E.

This confidence interval was used to generate model predictions of pharmacological perturbations. The pharmacological perturbations were: 1) sensory stimulation during application of the NPY receptor Y1 antagonist BIBP (Fig. S2F, red shaded area); 2) OG stimulation of GABAergic interneurons during application of BIBP (Fig. S2G, red shaded area), and, 3) OG stimulation of pyramidal neurons during inhibition of glutamatergic signaling by the administration of the AMPA and NMDA receptor antagonists CNQX and AP5, respectively (Fig. S2H, red shaded area). Correspondingly, experimental data of these perturbations are found in Uhlirova *et al*. [2], and shown as black symbols in Fig.S2F–H. As seen, the model predictions overlap with experimental data, qualitatively correctly predicting the dynamics observed in the experimental data.

**Figure S2.**
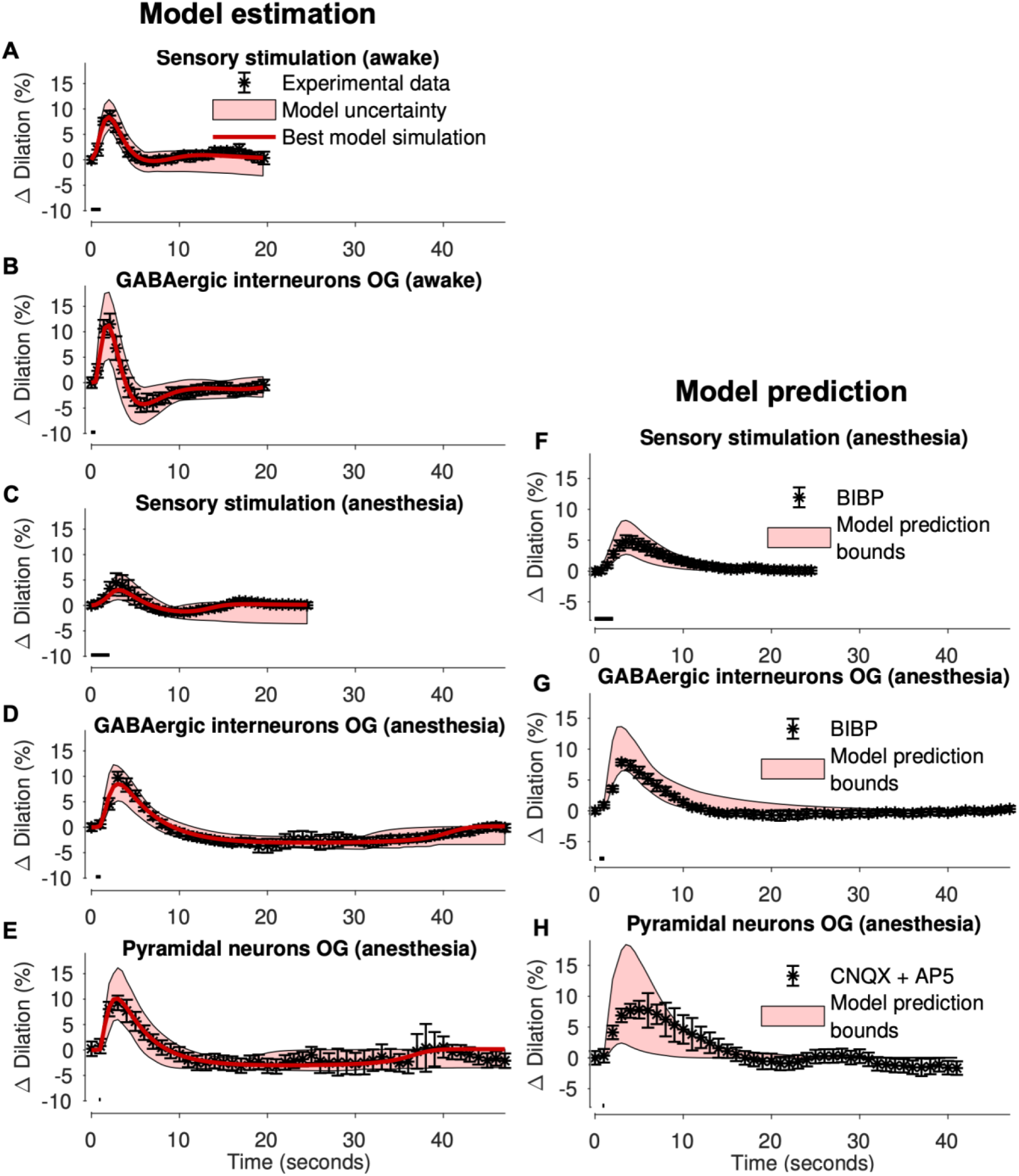
Model evaluation to arteriolar response data from awake (A–B) and anesthetized animals (C–H) for both optogenetic (OG) (B, D–E, G–H) and sensory (A, C, F) stimuli. The model parameters were fitted to data from A–E, while F–H was used as validation data. **A–E:** Model estimation paired with experimental data. For each graph, best estimated model simulation (solid red line) paired with model uncertainty (red shaded areas) compared to experimental data (black symbols, error bars depicting standard error of the mean). The stimulation length is indicated by the black bar in the lower-left portion of each graph. **F–H:** For each graph, model prediction bounds (red shaded areas) compared to experimental data (black symbols, error bars depicting standard error of the mean). The stimulation length is indicated by the black bar in the lower-left portion of each graph. Experimental data were extracted from Uhlirova *et al*. [3].

### S2 Posterior probability profiles of the model parameters

#### S2.1 Posterior probability profiles for model estimation in Figure 3

**Figure S3.**
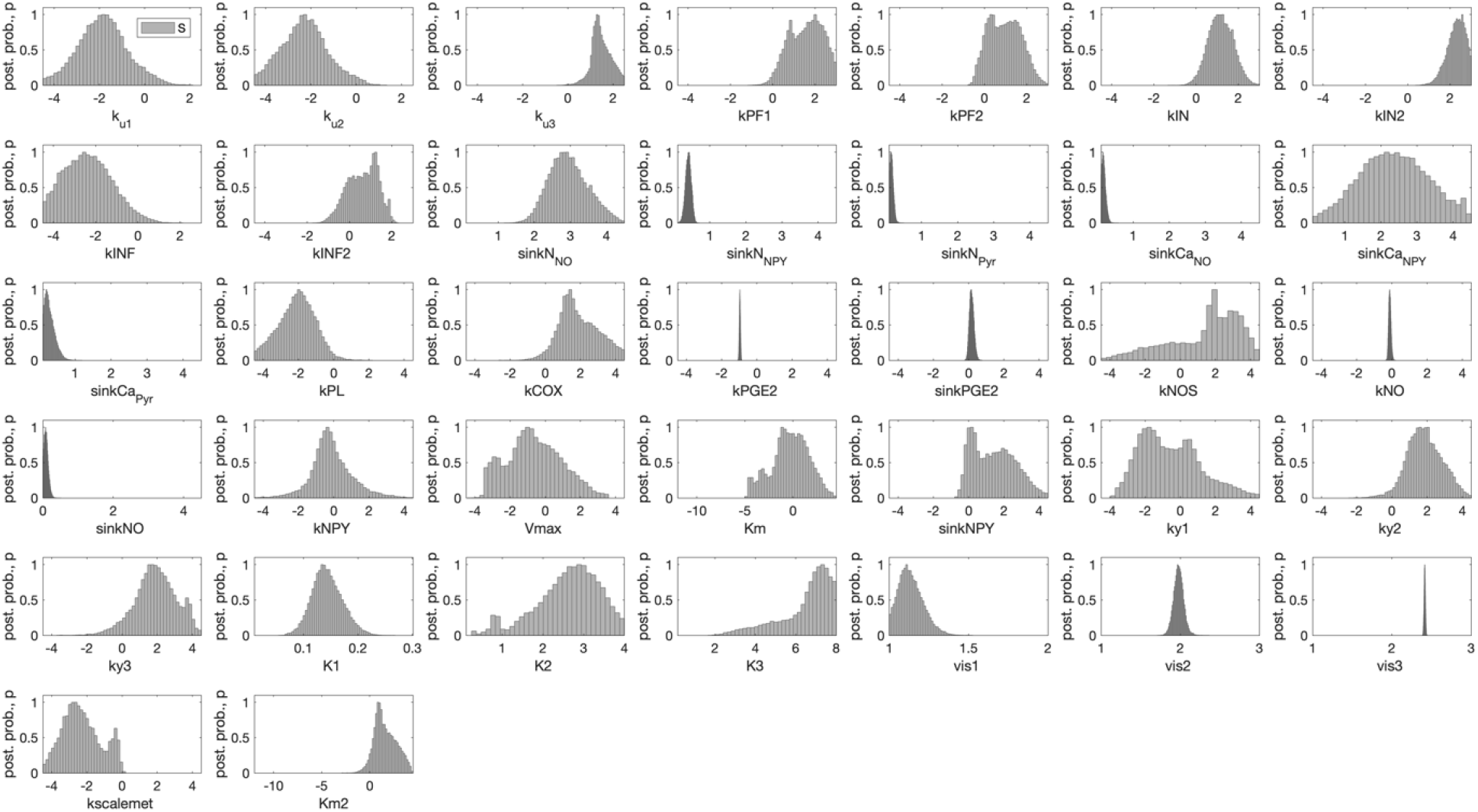
Posterior probability profile (y-axis) for each estimated model parameter (x-axis, log_10_ space) for the model estimated to data from Drew *et al*. [1].

**Figure S4.**
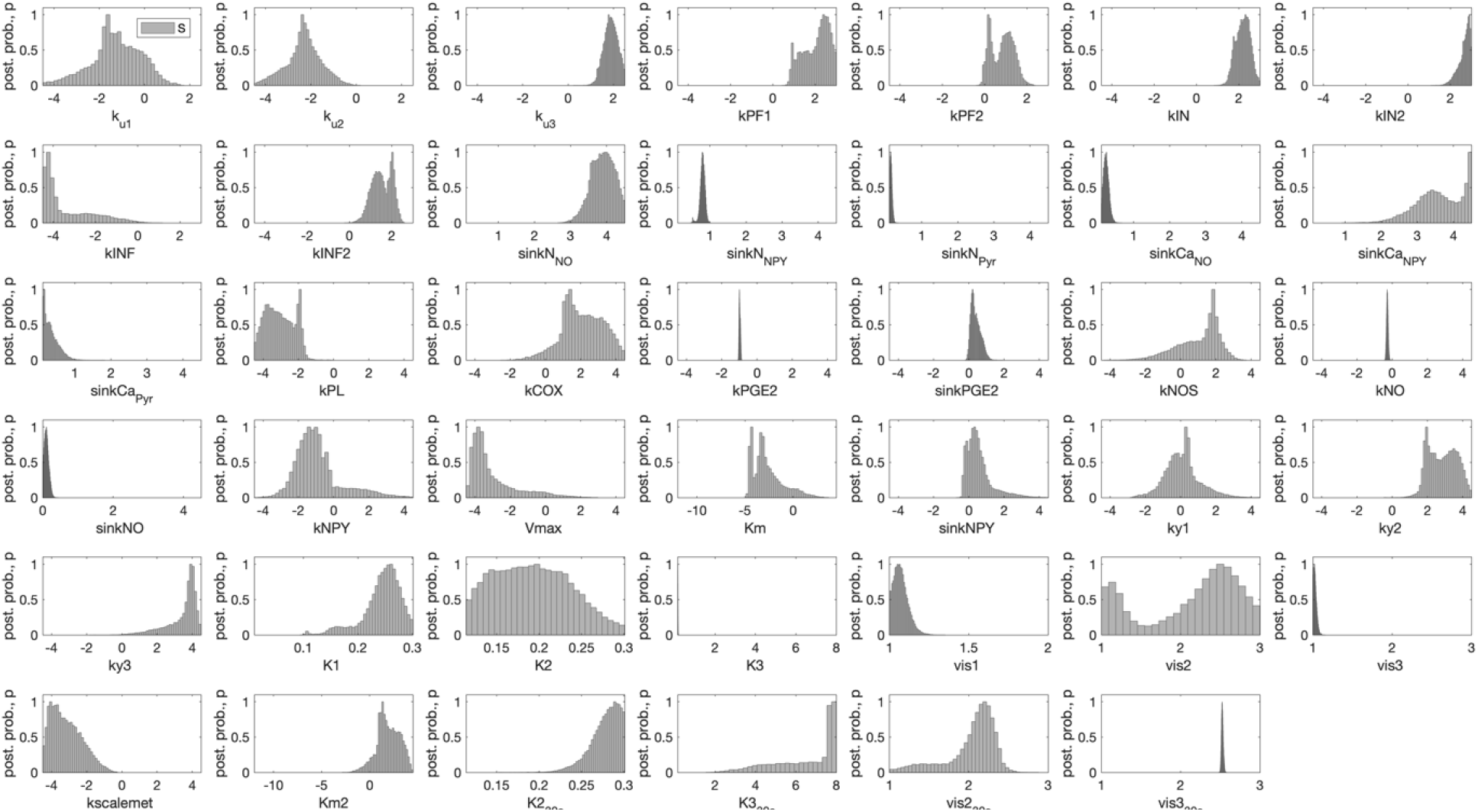
Posterior probability profile (y-axis) for each estimated model parameter (x-axis, log_10_ space) for the model estimated to data from Drew *et al*. [1], allowing the viscoelasticity and stiffness coefficients of the capillary and venular compartment change between 10 and 30 s stimulation.

#### S2.2 Posterior probability profiles for model estimation in Figure S2

**Figure S5.**
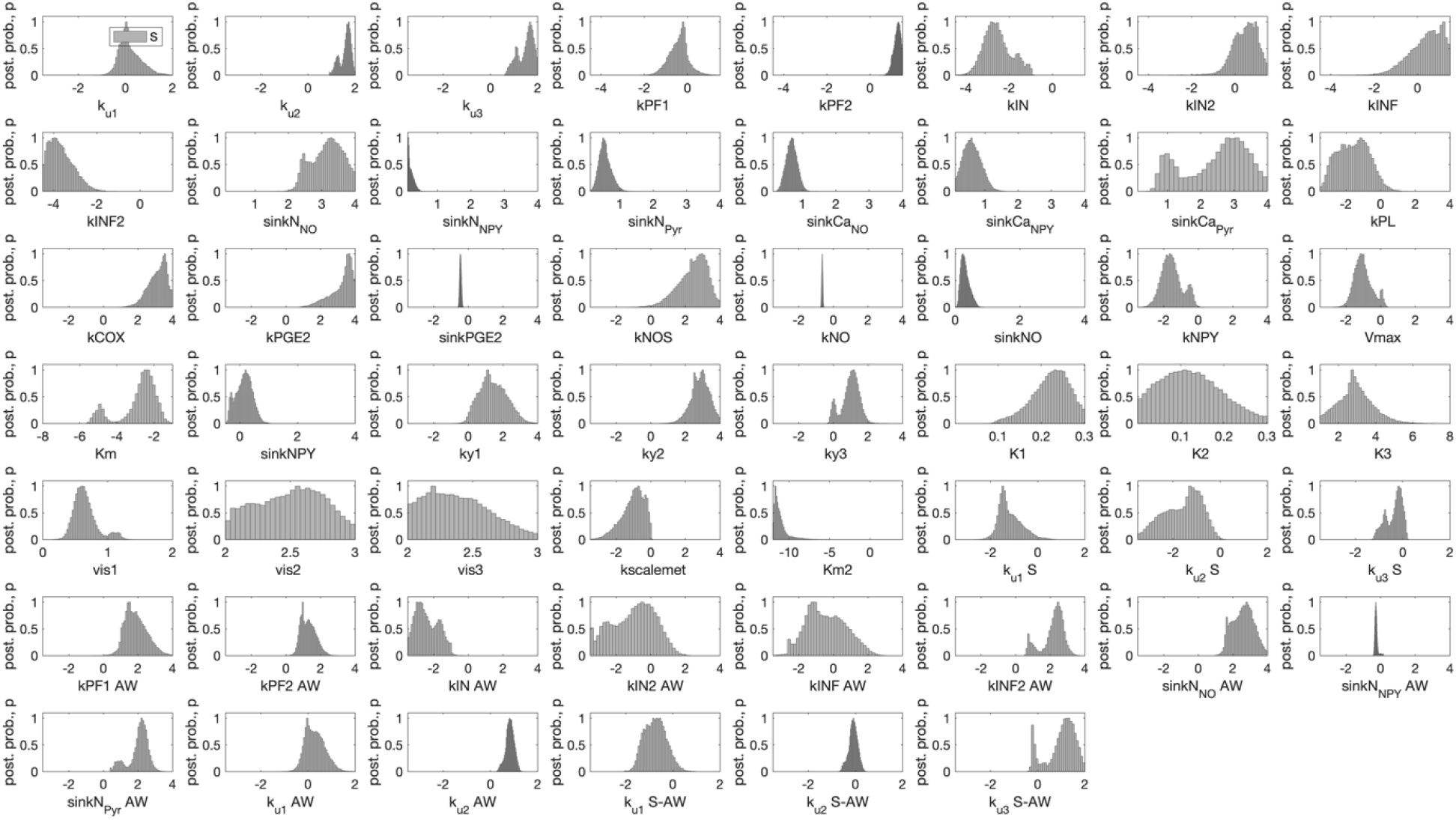
Posterior probability profile (y-axis) for each estimated model parameter (x-axis, log_10_ space) for the model estimated to data from Uhlirova *et al*. [2]. Two affixes are used: ‘AW’ which represent parameter value during awake condition, and ‘S’ which represent value for sensory stimulation.

#### S2.3 Posterior probability profiles for model estimation in Figure 5

**Figure S6.**
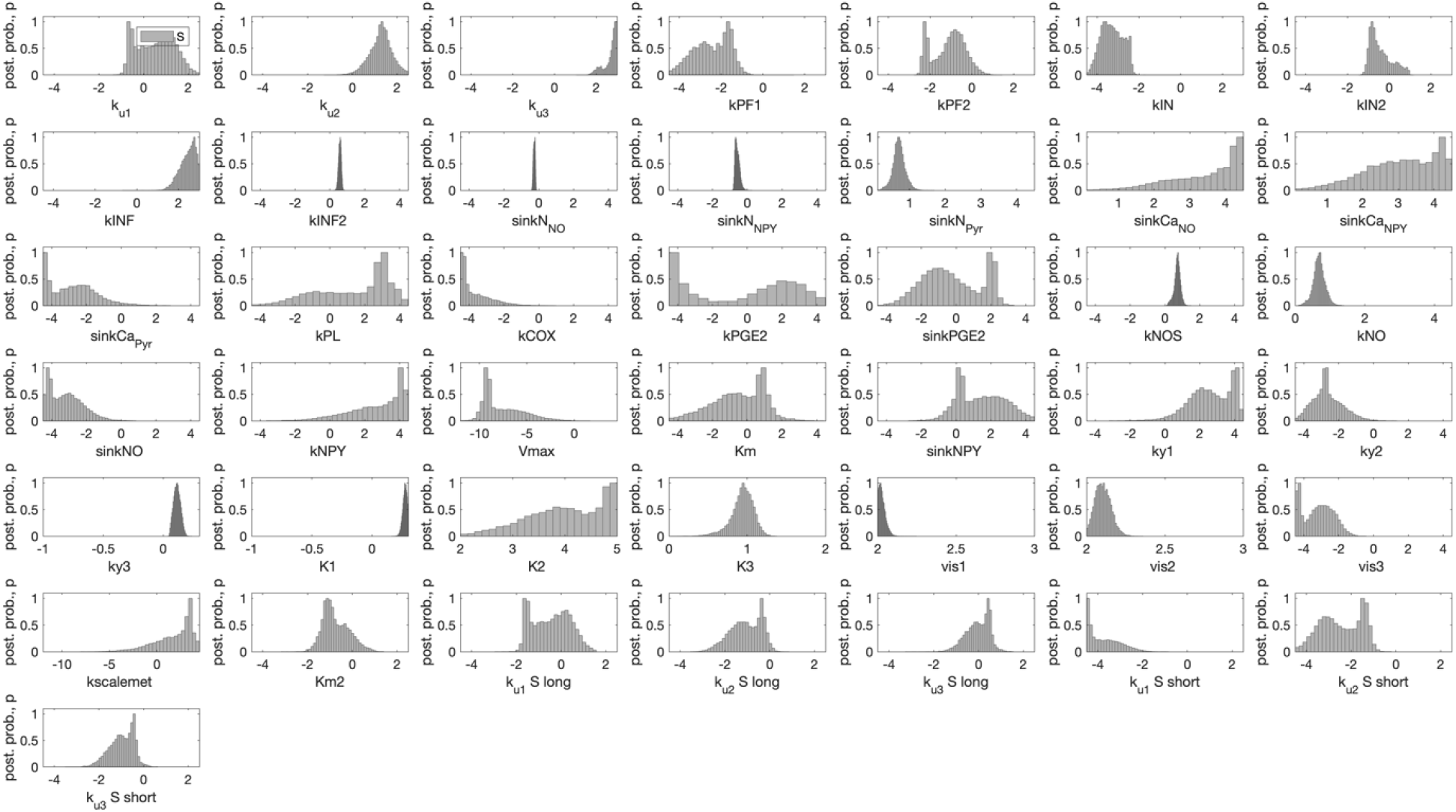
Posterior probability profile (y-axis) for each estimated model parameter (x-axis, log_10_ space) for the model estimated to data from Desjardins et al. [4]. The affix ‘S’ represent parameter value for sensory stimulation and is further subdivided into short and long sensory stimulation.

#### S2.4 Posterior probability profiles for model estimation in Figure 6

**Figure S7.**
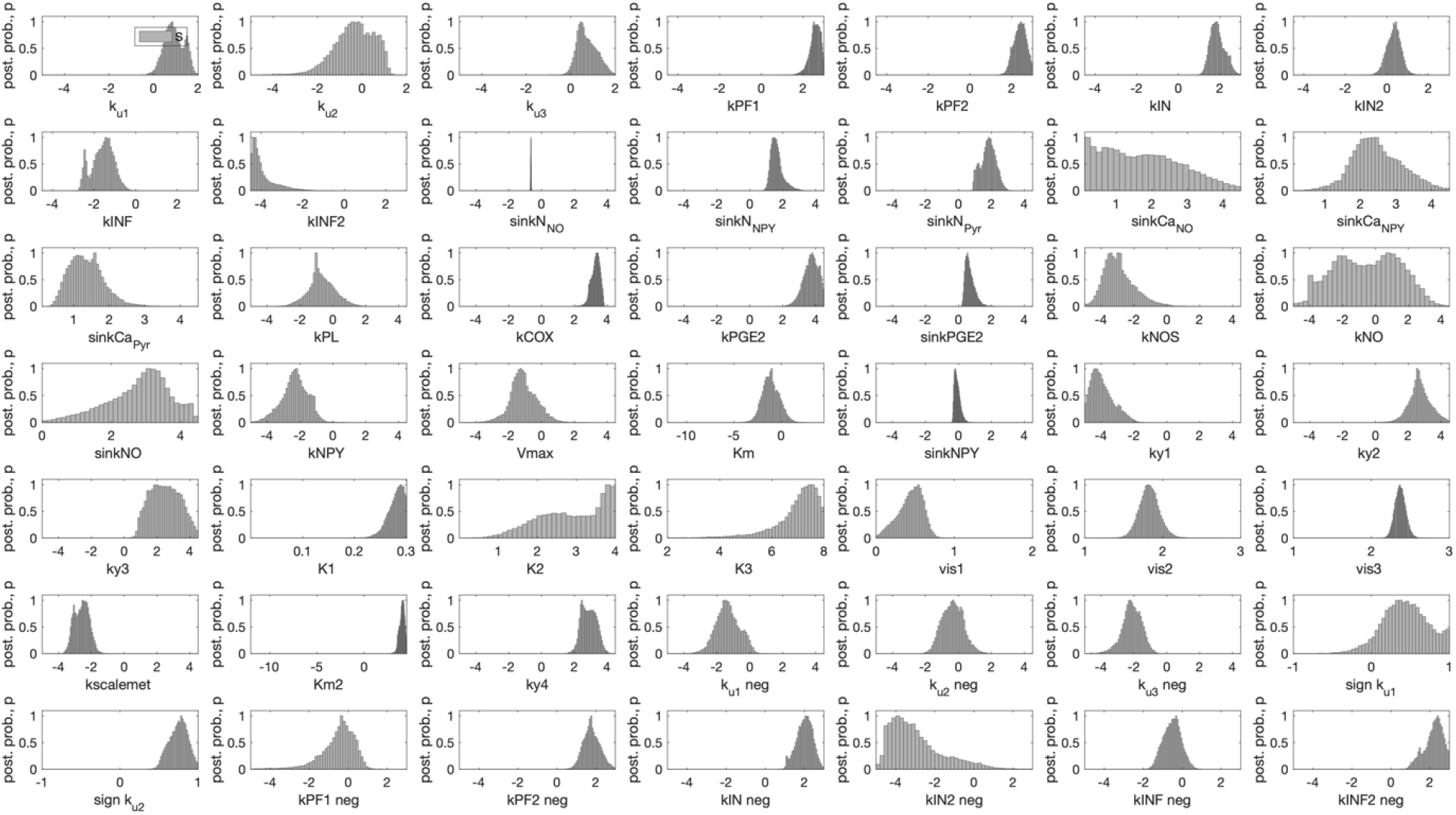
Posterior probability profile (y-axis) for each estimated model parameter (x-axis, log_10_ space) for the model estimated to data from Shmuel et al. [5]. The affix ‘neg’ represent parameter value for negative response.

#### S2.5 Posterior probability profiles for model estimation in Figure 7

**Figure S8.**
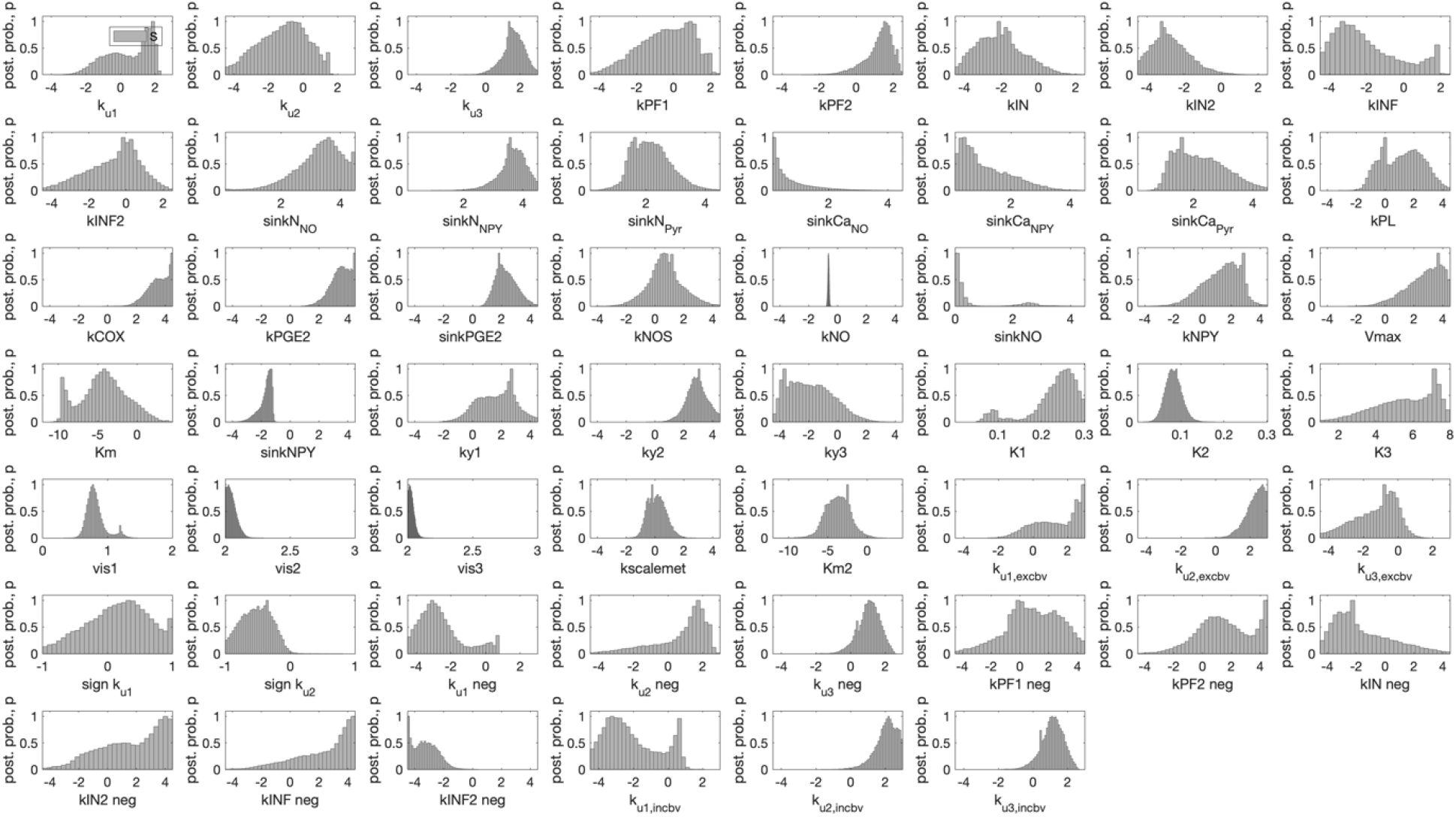
Posterior probability profile (y-axis) for each estimated model parameter (x-axis, log_10_ space) for the model estimated to data from Huber et al. [6]. The affix ‘neg’ represent parameter value for negative response. The affix ‘excbv’ and ‘incbv’ represent parameter values for the excitatory and inhibitory task, as presented in Figure 7BC, respectively.

